# Colocalized, Bidirectional Optogenetic Modulations in Freely Behaving Mice with a Wireless Dual-Color Optoelectronic Probe

**DOI:** 10.1101/2021.06.02.446749

**Authors:** Lizhu Li, Lihui Lu, Yuqi Ren, Guo Tang, Yu Zhao, Xue Cai, Zhao Shi, He Ding, Changbo Liu, Dali Cheng, Yang Xie, Huachun Wang, Xin Fu, Lan Yin, Minmin Luo, Xing Sheng

## Abstract

The precise control of neural activities at both cellular and circuit levels reveals significant impacts on the fundamental neuroscience explorations and medical applications. Optogenetic methods provide efficient cell-specific modulations, and the ability of simultaneous neural activation and inhibition in the same brain region of freely moving animals is highly desirable and being actively researched. Here we report bidirectional neuronal activity manipulation accomplished by a wireless, dual-color optogenetic probe in synergy with the co-expression of two spectrally distinct opsins (ChrimsonR and stGtACR2) in a rodent model. Based on vertically assembled, thin-film microscale light-emitting diodes (micro-LEDs) with a lateral dimension of 125 × 180 µm^2^ on flexible substrates, the dual-color probe shows colocalized red and blue emissions and allows chronic *in vivo* operations with desirable biocompatibilities. In addition, we discover that neurons co-expressing the two opsins can be deterministically evoked or silenced under red or blue irradiations. Implanted in behaving mice, the wirelessly controlled dual-color probe interferes with dopaminergic neurons in the ventral tegmental area (VTA), increasing or decreasing dopamine levels with colocalized red and blue stimulations. Such bidirectional regulations further generate rewarding and aversive behaviors of freely moving mice in a place preference test and interrogate social interactions among multiple mice. The technologies established here will create numerous opportunities and profound implications for brain research.

## INTRODUCTION

Understanding brain functions and treating neurological disorders rely on the continuous development of advanced technologies to interrogate complex nervous systems^1, 2^. Over the past decades, optogenetic methods have been emerging as a powerful toolset for effective and precise neural manipulation, owing to their capability of specific cell targeting^3, 4^. Until now, a range of genetically encoded, light-sensitive proteins (opsins) with distinct spectral responses have been engineered to regulate flows of different cations and anions^5–15^, triggering various biological responses, from cells and *C. elegans* to rodents and non-human primates^16–19^. Accompanying the development of opsins, advanced technologies for light delivery into the brain have also been progressing rapidly. Apart from various waveguide-based emitters interfacing with external light sources^20–22^, recently developed implantable probes based on thin-film, microscale optoelectronic devices have offered a viable solution to versatile neural modulations in untethered animals, when incorporated with various wirelessly operating systems based on radio frequency (RF) antennas^23–25^, near-field communication (NFC)^26^, Bluetooth chips^27^, and infrared receivers^28^. In addition to these achievements, bidirectional neural modulations, specifically, by activating or suppressing the same or different populations of cells in behaving animals with the light of different wavelengths, are highly demanded. Such capabilities would enable more precise control of neural activities to advance both neuroscience explorations and disease medications^29^. Certain efforts have been attempted, including explorations based on experiments by expressing excitatory and inhibitory opsins in separate animals^30, 31^ or distinct cells in different neural regions^17, 32^. There are also reports on dual-color optogenetic activation and inhibition by co-expressing different opsins in the same cells^33–36^. Recent efforts also remarkably demonstrate bidirectional optogenetic modulations with implantable waveguides coupled to extra-cranial dual-color laser sources and their combination with *in vivo* electrophysiological recordings^23, 36, 37^, but the wireless operation has not been achieved. Therefore, we envision that wirelessly operated light sources with polychromatic emissions, which collaborate with co-expressed opsins, can be implemented for untethered use in freely moving animals and enable previously inaccessible applications.

This study presents synergic optoelectronic and biological strategies to overcome challenges of existing techniques, and realize colocalized, bidirectional neural manipulations in untethered behaving mice. With heterogeneously integrated thin-film, microscale light-emitting diodes (micro-LEDs), a wirelessly operated, implantable dual-color probe allows independently controlled red and blue emissions in the same region. We also discover that the co-expression of two channelrhodopsins (ChrimsonR and stGtACR2) in the same group of neurons enables efficient light-evoked cell depolarization and hyperpolarization with controlled cation and anion flows under red and blue illuminations, respectively. With the wireless, dual-color micro-LED probes implanted into the ventral tegmental area (VTA) of freely moving mice, preference and aversion tests of individual and social behaviors clearly demonstrate the bidirectional optogenetic excitation and inhibition capability. Furthermore, *in vivo* behavioral results associated with the red and blue stimulations in the VTA are validated by increased or decreased dopamine levels detected in the nucleus accumbens (NAc) in real-time.

## RESULTS

### Wirelessly operated, dual-color micro-LED probes and circuit systems

We design and implement a wireless, dual-color micro-LED probe that generates red and blue emissions in the same brain region for bidirectional optogenetic modulations *in vivo*, as schematically illustrated in Figure 1a. Our previous works establish wirelessly operated microscale optoelectronic device strategy for *in vivo*, single-color optogenetic stimulations^37–40^, here these concepts are applied for dual-color, bidirectional modulations. Specifically, with the co-expression of spectrally distinct ion channels, ChrimsonR^7^ and stGtACR2^9^, in the same population of cells, red and blue light regulates the inflow of cations and anions, causing cell depolarization and hyperpolarization, respectively (Figure 1b). Figure 1c displays an enlarged view of the dual-color probe, with design and fabrication details provided in the Methods and Figure S1. From bottom to top, the probe structure comprises of a copper (Cu) coated polyimide (PI) thin-film substrate, an indium gallium phosphide (InGaP) red LED, a silicon oxide (SiO_2_) / titanium oxide (TiO_2_) based dielectric filter, and an indium gallium nitride (InGaN) blue LED. The Cu coating on PI serves as an efficient heat spreader to reduce the probe’s operation temperature^37^. All the devices (red LEDs, blue LEDs, and filters) are formed in thin-film, microscale formats (lateral dimension ~ 125 µm × 180 µm, thickness ~ 5–7 µm) through epitaxial liftoff, and vertically assembled via transfer printing^37, 39, 40^ (Figure 1d). The dimensions of our LEDs are designed to target a large nucleus like VTA. Although smaller devices down to 10–20 µm can be fabricated^41, 42^, reducing LED size decreases luminescence efficiencies^43, 44^ and may cause additional tissue heating. The intrinsic optical transparency of active layers in the InGaN blue micro-LED allows emissions from the red micro-LED to pass through with minimal losses. Additionally, the designed filter selectively reflects blue light and transmits red light, enhancing the emissive efficiencies of both micro-LEDs.

**Figure 1.**
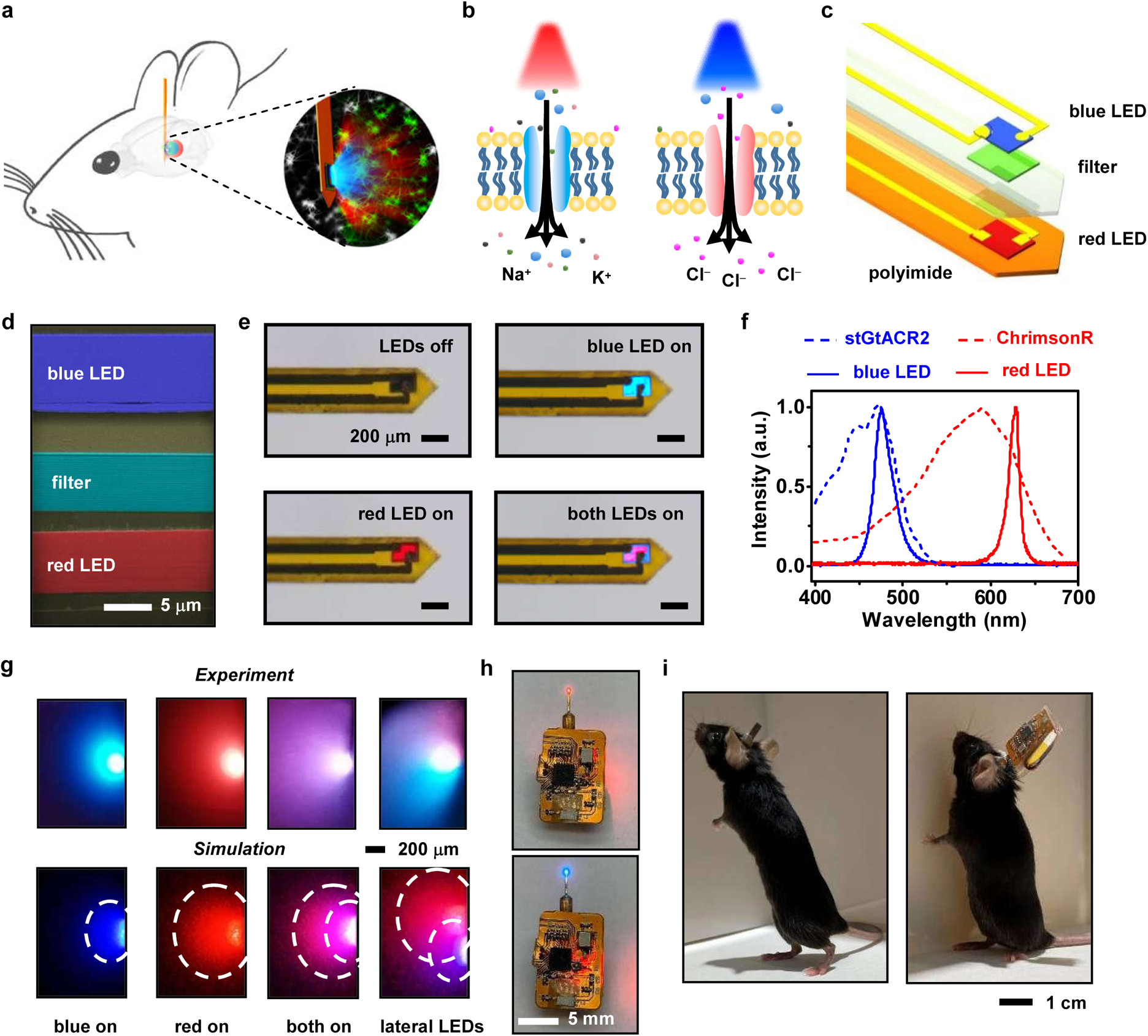
Wirelessly operated, dual-color microscale LED (micro-LED) probes for bidirectional, colocalized optogenetic control of neural activities. **a**, Schematic view of the probe inserted into the target brain area to perform dual-color neural activation and inhibition. **b**, Schematic, cellular scale depiction of the modulation function: red light regulates the cation channel ChrimsonR for depolarization, while blue light regulates the anion channel stGtACR2 for hyperpolarization. **c**, Exploded view of the dual-color LED probe made from vertically stacked blue and red thin-film micro-LEDs assembled on a flexible polyimide (PI) substrate, with a thin-film filter interface for spectrally selective reflection and transmission. **d**, Cross-sectional scanning electron microscopic (SEM) image of the probe structure, with false colors highlighting (from bottom to top) the indium gallium arsenide (InGaP) red micro-LED, the multilayer silicon dioxide / titanium dioxide (SiO_2_/TiO_2_) filter, and the indium gallium nitride (InGaN) blue micro-LED. Layers of SU-8 photoresists serve as the bonding materials. **e**, Optical images of a micro-LED probe (with a thickness of ~120 µm and a width of ~320 µm), showing independent control of blue and red emissions under current injections. **f**, Emission spectra of the blue and red micro-LEDs, in comparison with the action spectra of ion channels stGtACR2 and ChrimsonR. **g**, Experimental photographs and simulated results illustrating blue and red light propagations for a micro-LED probe embedded into a brain phantom (column 1–3). Results for a probe with laterally assemble blue and red LEDs are also presented (column 4), showing misplaced blue and red emissions. Iso-intensity lines show 10% of the maximum power. **h**, Images of a micro-LED probe integrated with a wireless circuit for independent control of dual-color emissions. **i**, Photos of a behaving mouse after intracranial implantation of a probe, with (right) and without (left) the wireless circuit module.

Flexible PI substrates with vertically stacked micro-LEDs are laser milled and encapsulated with a waterproof coating, forming a needle-shaped probe with a dimension (thickness ~120 µm and width ~320 µm) similar to a conventional silica fiber (diameter ~220 µm) used in optogenetic stimulations. The Cu coated PI substrate has a measured Young’s modulus of ~15 GPa, softer than silicon (~180 GPa) and tungsten (~400 GPa), but still much harder than the brain tissue (~1 kPa). The formed probe has a bending stiffness of ~9 × 10^4^ pNm^2^, similar to metal electrodes used for electrophysiological studies^45^. Separate metallization and electrical insulation ensure that red and blue micro-LEDs can be independently lighted up and display spectrally varied illuminations (red, blue, or combined) in the same location (Figure 1e and Movie S1). Figure 1f plots measured electroluminescence (EL) spectra. The red and blue micro-LEDs exhibit emission peaks at 630 nm and 480 nm, which are in good accordance with the excitation spectra of ChrimsonR and stGtACR2 (data extracted from references^7, 8^), respectively. Moreover, the spectral separation between the two opsins, guarantee precise optogenetic activation and inhibition with minimal optical interferences. More optoelectronic properties of the dual-color micro-LED probe, including current-voltage characteristics, quantum efficiencies, power irradiance, and angular-dependent emissive profiles are presented in Figure S2. Under injection currents of 10 mA, the red and the blue micro-LEDs can reach irradiances up to 50 mW/mm^2^ and 200 mW/mm^2^ on the device surface, which are sufficient for optogenetic modulations. In addition, both devices exhibit a uniform, near Lambertian emission profile.

To illustrate the optical performance of the devices, we insert a dual-color probe into a brain phantom (Figure 1g, top). For comparison, established models calculate optical profiles based on the optical properties (absorption and scattering) of brain tissues (Figure 1g, bottom). Both experimental and simulation results reveal that vertically stacked dual-color micro-LEDs show colocalized red and blue emission profiles, ensuring effective activation and inhibition in the same brain region. Power irradiances for blue and red emissions decrease to 10% of the original power at propagating distances of ~400 µm and ~800 µm, respectively. Compared to a probe with laterally arranged red and blue micro-LEDs displaying misaligned emissions with a relatively small overlapping volume of light, the vertically assembled device provides a smaller footprint and generates an optimal overlapping profile for dual-color stimulations.

Thermal behaviors of the micro-LED probe are also systematically analyzed, with detailed results presented in Figures S3 and S4. Probes are inserted into the brain phantom as well as the living mouse brain, with temperature rises measured by an infrared camera and a thermocouple. Both experiments obtain similar results and are in agreement with the thermal models established based on finite-element analysis. Under a pulsed operation (20-Hz frequency, 10-ms pulse duration, current 0–10 mA), the tissue temperature rises caused by the red micro-LED can be restricted within 1 °C. The operation of the blue micro-LED induces a higher thermal effect because practical neural inhibition for stGtACR2 requires continuous current injection; nonetheless, the tissue heating can be controlled within 2 °C by limiting the current within 5 mA. Such operation modes for red and blue micro-LEDs will be employed throughout *in vivo* experiments, to mitigate undesired neural responses and possible tissue damage due to overheating.

We also design a customized, miniaturized flexible circuit module to wirelessly operate the dual-color micro-LED probe. Compared to previous reports^22, 36^ on tethered fiber or laser coupled devices for dual-color stimulations, the wireless operation here enables the study of complex behaviors of freely moving animals more conveniently. Controlled by radio frequency (RF) communication at 2.4 GHz, the battery-powered circuit (weight ~1.9 g) independently addresses the red and the blue micro-LEDs, by adjusting their pulse frequencies, durations, and injection currents for versatile neural modulations *in vivo* (Figure 1h, and details in the Methods and Figures S5–S9). Designed programming commands also allow the independent operation of multiple micro-LED probes in synchronized or non-synchronized modes (Movie S2). After subcranial implanting the micro-LED probe into a behaving mouse, the light-weighted circuit mounted onto the mouse head provides the capability for optogenetic control in an untethered manner (Figure 1i).

### Bidirectional optogenetic manipulations of neurons *in vitro*, with dual-color stimulations

We first evaluate the feasibility of dual-color, bidirectional optogenetic modulations for neurons co-expressing ChrimsonR and stGtACR2 in brain slices with electrophysiological tests. Mixed vectors of AAV-CAG-DIO-ChrimsonR-mCherry and AAV-EF1a-DIO-stGtACR2-EGFP are injected into the ventral tegmental area (VTA) of *DAT-Cre* mice (Figure 2a). The deployment of the soma-targeted variant stGtACR2 exhibits performance superior to previously reported anion channel GtACR2, by enhancing the membrane trafficking and reducing the axonal excitation^8, 9^. Figure 2b shows the fluorescence images of stained cells 3 weeks after viral injection in the VTA, including 4’,6-diamidino-2-phenylindole (DAPI, blue), stGtACR2 (green), and ChrimsonR (red). The merged graph and zoomed-in views (Figure 2c) clearly reveal the desirable co-expression of the two opsins in target cells. Statistical results show that ~30% of cells are labeled with stGtACR2 or ChrimsonR, and about 83% of these labeled cells co-express both opsins on average (Figure 2d).

**Figure 2.**
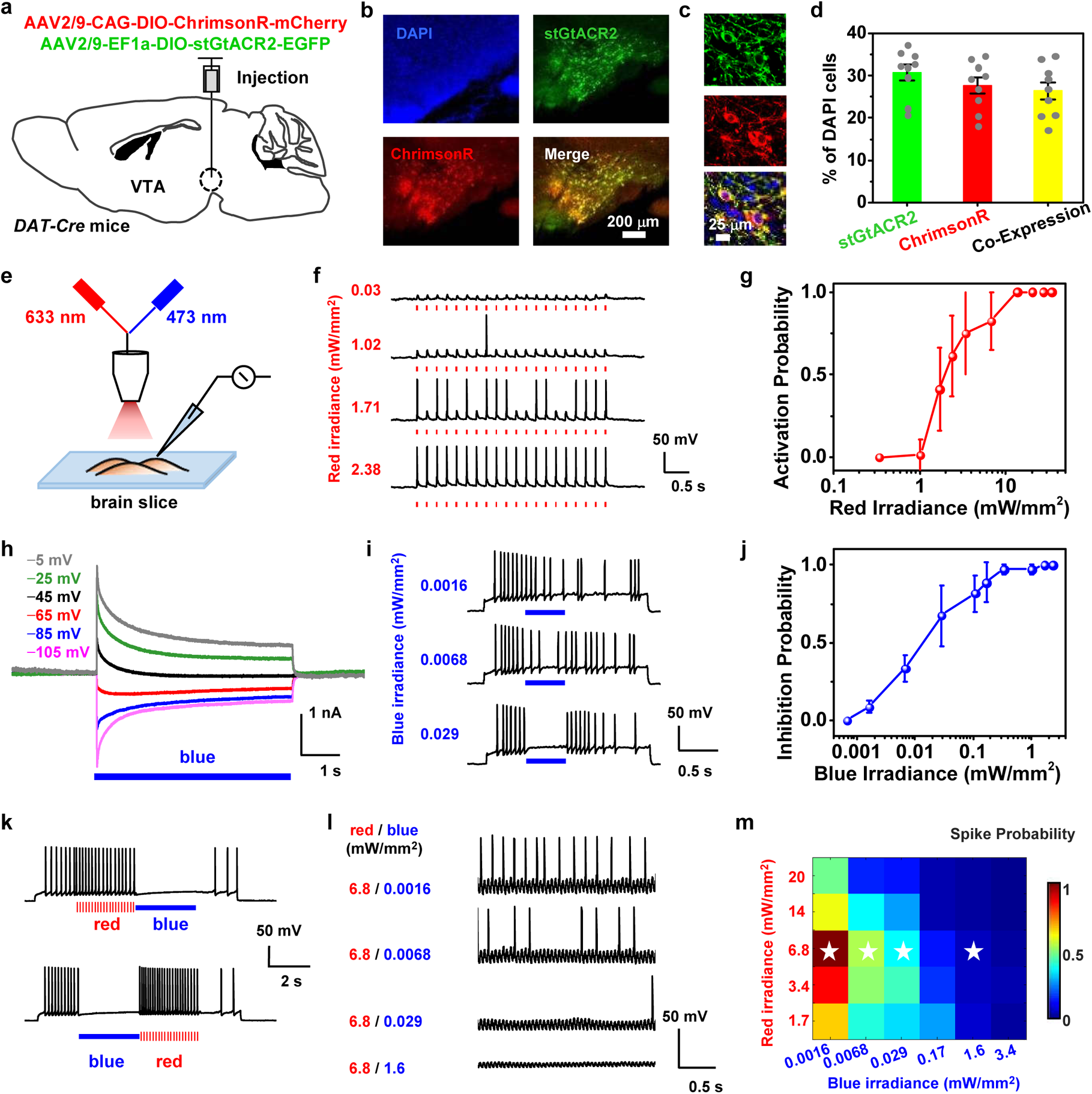
Bidirectional optogenetic activation and inhibition of neurons with red and blue light stimulation. **a**, Schematic strategy for co-expressing ChrimsonR and stGtACR2 in the ventral tegmental area (VTA) of *DAT-cre* mice. **b**, Immunostained fluorescence images of the VTA region expressing DAPI (blue), stGtACR2 (green), and ChrimsonR (red) and a merged image, 3 weeks after viral injection. **c**, Zoomed view showing cells co-expressing stGtACR2 (green) and ChrimsonR (red). **d**, Percentages of stGtACR2-, ChrimsonR- and co-expressing neurons among DAPI cells in the VTA (mean ± s.e.m., 3 slices per mouse, *n* = 3 mice). **e**, Illustration of the experiment setup with combined blue (480 nm) and red (633 nm) laser beams (spot diameter ~1 mm) for optogenetic stimulating brain slices during patch-clamp recordings *in vitro*. **f**, Example traces showing neural activation by pulsed red light at different irradiances (20 Hz, 10-ms pulse, 20-pulse train). **g**, Activated spike probability as a function of red irradiances (mean ± s.e.m., *n* = 4 cells). **h**, Example photocurrents of neurons at membrane potentials varied from −105 mV to −5 mV (bottom to top), under continuous blue light illumination (duration 5 s, irradiance 8 mW/mm^2^). **i**, Example traces of neural inhibition by continuous blue light at different irradiances (duration 0.5 s). **j**, Inhibited spike probability as a function of blue irradiances (mean ± s.e.m., *n* = 5 cells). **k**, Traces presenting bidirectional spike activation and inhibition with alternating red and blue illuminations (Patching current +90 pA; Red: duration 3 s, 20 Hz, 10-ms pulse, 2.2 mW/mm^2^; Blue: duration 3 s, 8.5 mW/mm^2^). **l**, Example traces of neural activities with simultaneous red and blue illuminations at different irradiances (duration 6 s). **m**, Summarized plot of measured spike probability (normalized mean values) as a function of red and blue irradiances (*n* = 5 cells). The stars indicate the position of the four example traces shown in Fig. 2l.

Whole-cell patch recordings capture intracellular signals of these cells under optical stimulations (Figure 2e). Other than using our designed micro-LED probe, here we employ a collimated light spot with coupled red (633 nm) and blue (480 nm) laser sources for *in vitro* experiments, to achieve uniform illuminations for better power calibrations. Figures 2f and 2g validate that pulsed red stimulations (20-Hz frequency, 10-ms pulse duration) induce action potentials (APs or spikes) by depolarizing these cells, and spike activation probability increases at elevated red irradiances. In these same cells, continuous blue stimulations suppress activities by hyperpolarization (Figures 2h–2j). Voltage clamp results illustrate that blue light evokes photocurrent reversion when the membrane potential becomes more positive (Figure 2h), and the probability for spike inhibition also increases at elevated blue irradiances (Figures 2i and 2j). Threshold irradiances for complete spike activation and inhibition are around 10 mW/mm^2^ and 1 mW/mm^2^ for red and blue illuminations, respectively. The results are consistent with previous reports on cells expressing only ChrimsonR or stGtACR2^7, 9^.

Notably, we do not only demonstrate that the co-expression of both opsins does not affect the efficacy of cell activity excitation or suppression, but also accomplish bidirectional activation and inhibition in the same cell by instantaneously altered red and blue irradiations (Figure 2k). In accordance with the activation spectra of two opsins (Figure 1f), red light does not interact with stGtACR2, which exhibits nearly zero responses at wavelengths longer than 600 nm. On the other hand, ChrimsonR should have responded to blue light, which could raise a concern about possible optical crosstalk. Nevertheless, spike activations caused by blue light do not occur in cells co-expressing ChrimsonR and stGtACR2, since cells are suppressed by turning on stGtACR2 channels at a much lower blue irradiance. This fact can be confirmed by patching the cells under combined red and blue illuminations (Figures 2l and 2m). Example traces in Figure 2l show that red-light-evoked spikes are suppressed under simultaneous blue illuminations even with very low irradiances. More traces are reported in Figure S10, and Figure 2m summarizes the spike probability (measured and averaged among *n* = 5 cells) with combined red and blue illuminations. Introducing blue irradiances orders of magnitude lower than red irradiances can effectively suppress cell spikes and cause activity inhibition.

### Bidirectional optogenetic modulation and simultaneous electrophysiological recording *in vivo*

Neural activities associated with dual-color stimulations are further examined with simultaneous electrophysiological recordings *in vivo* (Figure S11). Two opsins (ChrimsonR and stGtACR2) are co-expressed in the primary somatosensory cortex of *CaMKII-Cre* mice (Figure S11a). Followed by two weeks of recovery and viral expression, extracellular electrodes, and dual-color micro-LED probes are inserted in the head-fixed mice, with a separation of ~500 μm (Figure S11b). Spontaneous and red-light stimulated waveforms are recorded and sorted, showing typical extracellular spike events (Figure S11c). Multiple trials collected for a sample unit indicate that cell spike rates can be enhanced or decreased by applying red or blue illuminations (Figures S11d and S11e). Results summarized for multiple cells show that the basal spike firing rate (13 Hz) increases to 28 Hz during red illumination and decreases to 2 Hz during blue illumination (Figures S11f and S11g). These results showcase the capability for *in vivo* bidirectional modulations and are in good accordance with the previous report based on a different opsin combination (ChR2 and Jaws)^36^.

### Bidirectional optogenetic modulation of preference and aversion behaviors *in vivo*, with wireless dual-color micro-LED probes

Activities of the dopaminergic neurons in the VTA we manage to manipulate in Figure 2 significantly correlate with the reward and aversion-based behaviors^46–48^. In Figure 3, we further exploit the capability of bidirectional optogenetic manipulations by implanting dual-color micro-LED probes into behaving mice. After the injection of AAV-CAG-DIO-ChrimsonR-mCherry and AAV-EF1a-DIO-stGtACR2-EGFP vectors, the dual-color micro-LED probe is implanted into the VTA of *DAT-cre* mice via standard stereotaxic surgery (Figure 3a and Figure S12). Figure 3b presents fluorescence images of a brain slice with a micro-LED probe underneath it, showing the co-expression of ChrimsonR and stGtACR2 in the VTA, combined with the luminescence of the red and the blue micro-LEDs under corresponding optical excitations. To assess the devices’ biocompatibility *in vivo*, histological examinations of brain slices show that inflammatory reactions of the micro-LED probe are similar to a conventional silica fiber (diameter ~220 µm) 1 day or 3 weeks’ post-implantation (Figure S13), owing to its thin-film geometry, mechanical flexibility, and bio-friendly encapsulation. Also, mass spectrometric analysis shows that the brain tissue has a minimal accumulation of dissolved Cu element 5 weeks’ post-implantation (Figure S14). In addition, we perform dynamic calcium imaging on acute brain slices for AAV-hsyn-GCaMP6m expressing mice 2 weeks’ post-implantation (Figure S15). Cell activities can be effectively recorded in neurons ~100 μm away from the lesion region, in agreement with the histological analysis. Moreover, the parylene-based, waterproof encapsulation maintains the micro-LEDs’ chronic and stable operation for more than 200 days after implantation (Figure S16).

**Figure 3.**
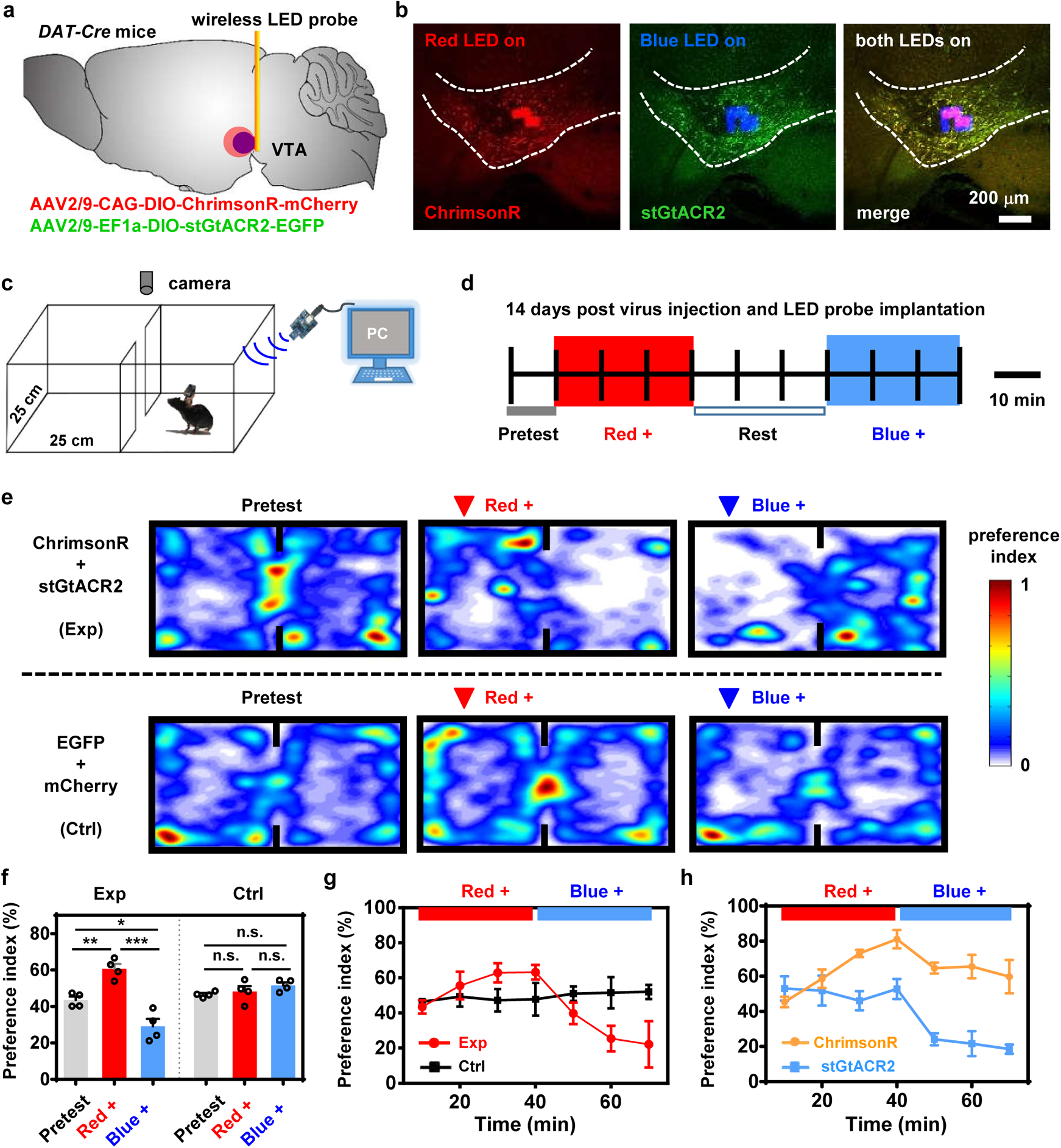
Bidirectional, *in vivo* optogenetic regulation of mice preference or aversion behaviors with wirelessly operated, dual-color microscale LED (micro-LED) probes. **a**, Diagram illustrating the combined viral injection (AAV2/9-CAG-DIO-ChrimsonR-mCherry and AAV2/9-EF1a-DIO-stGtACR2-EGFP) followed by implanting the micro-LED probe into the VTA of *DAT-cre* mice. **b**, Micrographs of the VTA showing co-expression of stGtACR2 and ChrimsonR, with a dual-color micro-LED probe embedded into the tissue. **c**, Illustration of the real-time place preference test with a two-compartment arena, where a mouse receives signals for dual-color control and its locomotion is recorded by a camera. **d**, Patterns used for optogenetic modulations, including a 10-min pretest, a 30-min red LED stimulation (20 Hz, 10-ms pulse, current 7 mA), a 30-min rest, and a 30-min blue LED stimulation (continuous, current 5 mA). **e**, Representative heat maps showing real-time preference and aversion behavior following red or blue stimulation for mice co-expressing stGtACR2 + ChrimsonR (experiment, or Exp) or EGFP + mCherry (control, or Ctrl). **f**, Summary of preference indices (the ratio of the time that mice spend in the left chamber to the whole recorded time) under red and blue stimulations for Exp and Ctrl groups (*n* = 4 mice each). Student’s *t* test, * *P* < 0.05, ** *P* < 0.01, *** *P* < 0.001, n.s. *P* > 0.05. **g**, Preference indices measured at different times under red and blue stimulations for Exp and Ctrl groups (*n* = 4 mice each). **h**, Preference indices measured at different times under red and blue stimulations for groups expressing only ChrimsonR or stGtACR2 (*n* = 3 mice each).

*In vivo* experiments are performed within a real-time place preference/avoidance (RTPP) paradigm. After 14 days of virus and micro-LED probe implantation, we place mice in a two-compartment apparatus and monitor their behaviors with a camera during optogenetic modulations (Figure 3c and Figure S17). Figure 3d provides the test protocol, including a baseline pretest (10 mins), red LED stimulation (pulsed wave, 30 mins), a rest phase (30 mins), and blue LED stimulation (continuous wave, 30 mins). The red or blue LED stimulations are only provided once the mice appear in the left chamber. Representative heat maps in Figure 3e show the change of real-time behaviors during optogenetic stimulations, for mice expressing different combinations of opsins. Summarized results in Figure 3f clearly show that mice co-expressing ChimsonR and stGtACR2 present strong preference and avoidance behaviors following red and blue stimulations, respectively. Additionally, the application of both red and blue stimulations in opposite chambers further enhances the preference indices of mice compared to those only experiencing single color stimulations as well as the control group (co-expressing EGFP and mCherry), demonstrating the efficacy of bidirectional modulation (Figure S18).

By contrast, the control groups without photosensitive opsins (expressing only EGFP and mCherry) show little difference of behaviors under optical stimulations. More results are provided in Figures S19 and S20, comparing these mice with other groups expressing only ChrimsonR or stGtACR2. When introducing only ChrimsonR, mice exhibit place preference under both red and blue stimulations. For the group with only stGtACR2, blue light effectively evokes place aversion while red light generates little influence on mouse behaviors. Figures 3g and 3h present the dynamic behavior change at different time courses. These results are consistent with the spectral characteristics of the opsins as well as the *in vitro* neural responses at the cellular level (Figure 2), elucidating the utility of bidirectional modulations with wireless, dual-color micro-LED probes. In conclusion, the implantable wireless dual-color LED probe combined with two spectrally distinct channelrhodopsins can realize bi-directional control of the place preference or aversion in freely behaving mice.

### Bidirectional optogenetic modulation of social interactions among multiple mice, with wireless dual-color micro-LED probes

Modulating dopaminergic neurons in the VTA region also influences the social preference, in particular, affective and aggressive behaviors^49, 50^. Compared with tethered stimulation systems, our wirelessly operated micro-LED probes with independently addressable illuminations present prominent advantages in studying social interactions among multiple objects (Movie S3). Additionally, the colocalized, dual-color emission capability allows the bidirectional regulation of social activities in real-time, which is difficult to access with single color stimulations^49, 51^. In paired-object interactions (Figure 4a), two mice co-expressing ChrimsonR and stGtACR2 in the VTA are placed in an open-field arena, and synchronized red or blue stimulations are provided for 5 mins each. Mice behaviors are video recorded throughout the test period, with social activities (including sniffing and attacking) labeled and scored (Figure 4b). The total time spent in these activities is summarized and compared among different scenarios (red light on, blue light on, and resting) in Figure 4c. In comparison to the resting state, red or blue LED stimulations strongly promote or suppress social interactions between two objects in the experiment group, while no significant effects are observed in the control group (expressing EGFP and mCherry).

**Figure 4.**
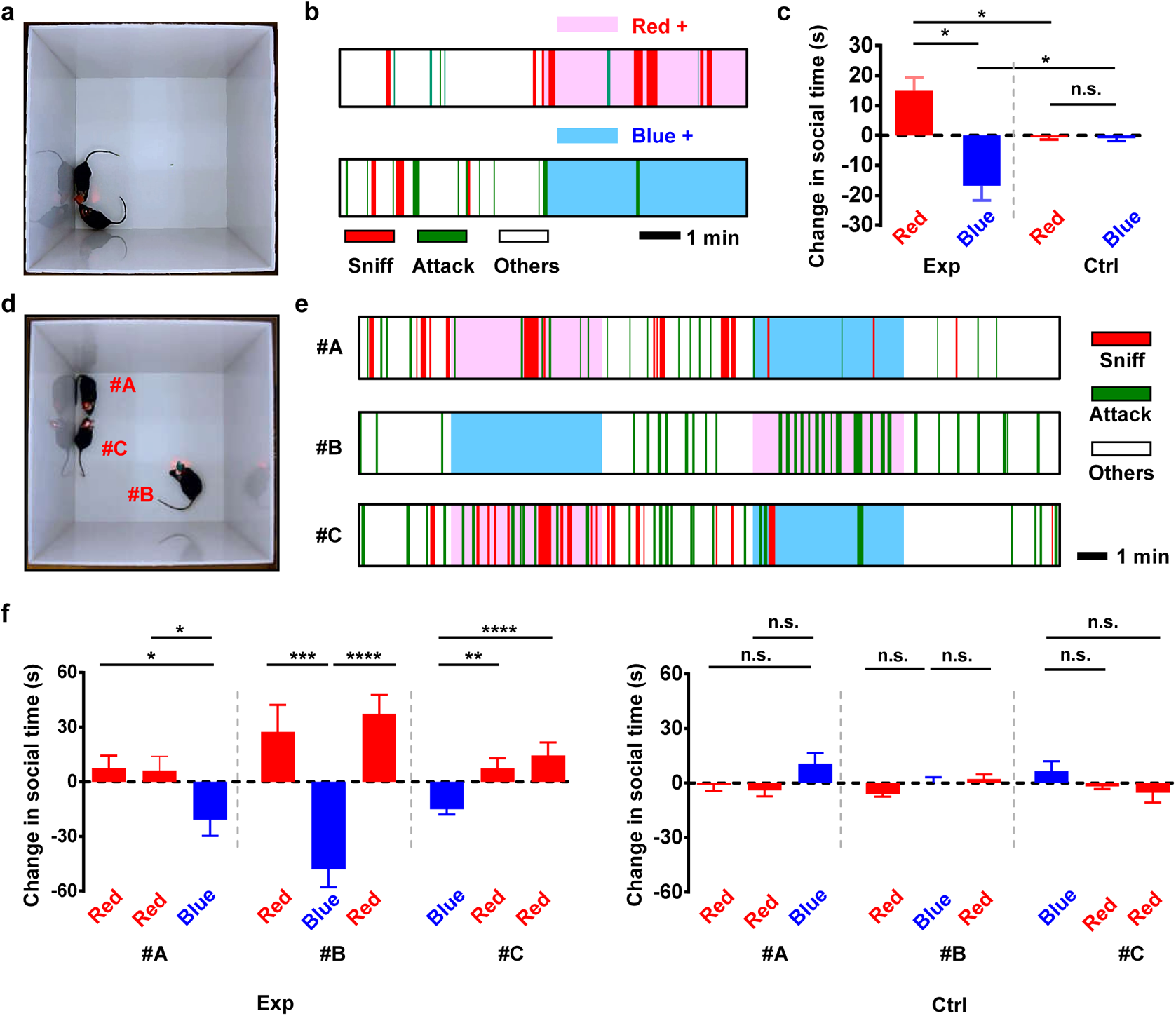
Bidirectional, *in vivo* optogenetic control of mice social interactions with wirelessly operated, dual-color micro-LED probes. **a**, Photograph of two *DAT-cre* mice with wireless probes in an arena (30 × 30 cm^2^). Experiment groups (Exp) express stGtACR2 + ChrimsonR, and control groups (Ctrl) express EGFP + mCherry. **b**, Representative behavioral sequences recorded for one mouse receiving red or blue stimulations during social interactions. Here sniffing and attacking behaviors are highlighted and labeled, and other behaviors include locomotion, resting and escaping. **c**, Summary of changes in social interactions under red or blue stimulations. Here we compare the time spent in sniff and attack during the 5 mins light stimulation with the 5 mins resting state. *n* = 4 groups for both Exp and Ctrl. **d**, Photograph of three *DAT-cre* mice with wireless probes in an arena. **e**, Representative behavioral sequences recorded for the three mice (#A, #B, #C) receiving red or blue stimulation during social interactions. **f**, Summary of changes in social interactions under red or blue stimulations for each mouse. *n* = 4 groups for both Exp (left) and Ctrl (right). All data are represented as mean ± s.e.m.. Student’s *t* test, * *P* < 0.05, ** *P* < 0.01, *** *P* < 0.001, **** *P* < 0.0001, n.s. *P* > 0.05.

The wireless micro-LED probes can further be applied to interrogate more complicated interaction behaviors among multiple (> 2) objects. In Figure 4d, we place three mice in the same social chamber and provide these animals with alternating red or blue LED stimulations. In such a testing paradigm, two mice receive red stimulations while the third one receives blue stimulation, then vice versa (Figure 4e). The mice with red stimulations spend more time interacting with each other compared to the individual under blue illuminations, and the social preference can be reversed by alternating the stimulation patterns among these three mice in real-time (Figure 4f). By contrast, summarized data among control groups present no significant difference under red or blue stimulations. Collectively, these social interaction explorations clearly demonstrate the unique advantage of our wireless and bidirectional LED stimulators in the interrogation of complex animal behaviors.

### Bidirectional optogenetic regulation of dopamine release *in vivo*, with wireless dual-color micro-LED probes

The effective bidirectional behavior manipulations demonstrated in Figures 3 and 4 can be ascribed to the light-induced activation and inhibition of dopaminergic neurons in the VTA. We further investigate the effect of optogenetic stimulations on dopamine release using *in vivo* experiments described in Figure 5a. Similarly, combined opsins (ChrimsonR and stGtACR2) are expressed and followed by implanting the dual-color micro-LED probe in the VTA of *DAT-cre* mice. While stimulations are performed in the VTA, dopamine signals are simultaneously monitored in one of its main projection areas, the nucleus accumbens (NAc)^52–54^. A genetically encoded indicator GRAB_DA2m_ is expressed in the NAc, where the fluorescence signals are recorded with a fiber photometer^30, 31, 48, 49, 56, 57^. The combination of micro-LED based optogenetic stimulations and fiber-based photometric recordings realizes all-optical interrogation of neural circuits. During the surgery, we carefully select the insertion direction of the micro-LED probe, to obtain desirable stimulation efficacy in the VTA and minimize the optical crosstalk that could disturb photometric recording in the NAc (Figure S21). Fluorescence images in Figure 5b show the expressions of GRAB_DA2m_ in the NAc as well as ChrimsonR and stGtACR2 in the VTA.

**Figure 5.**
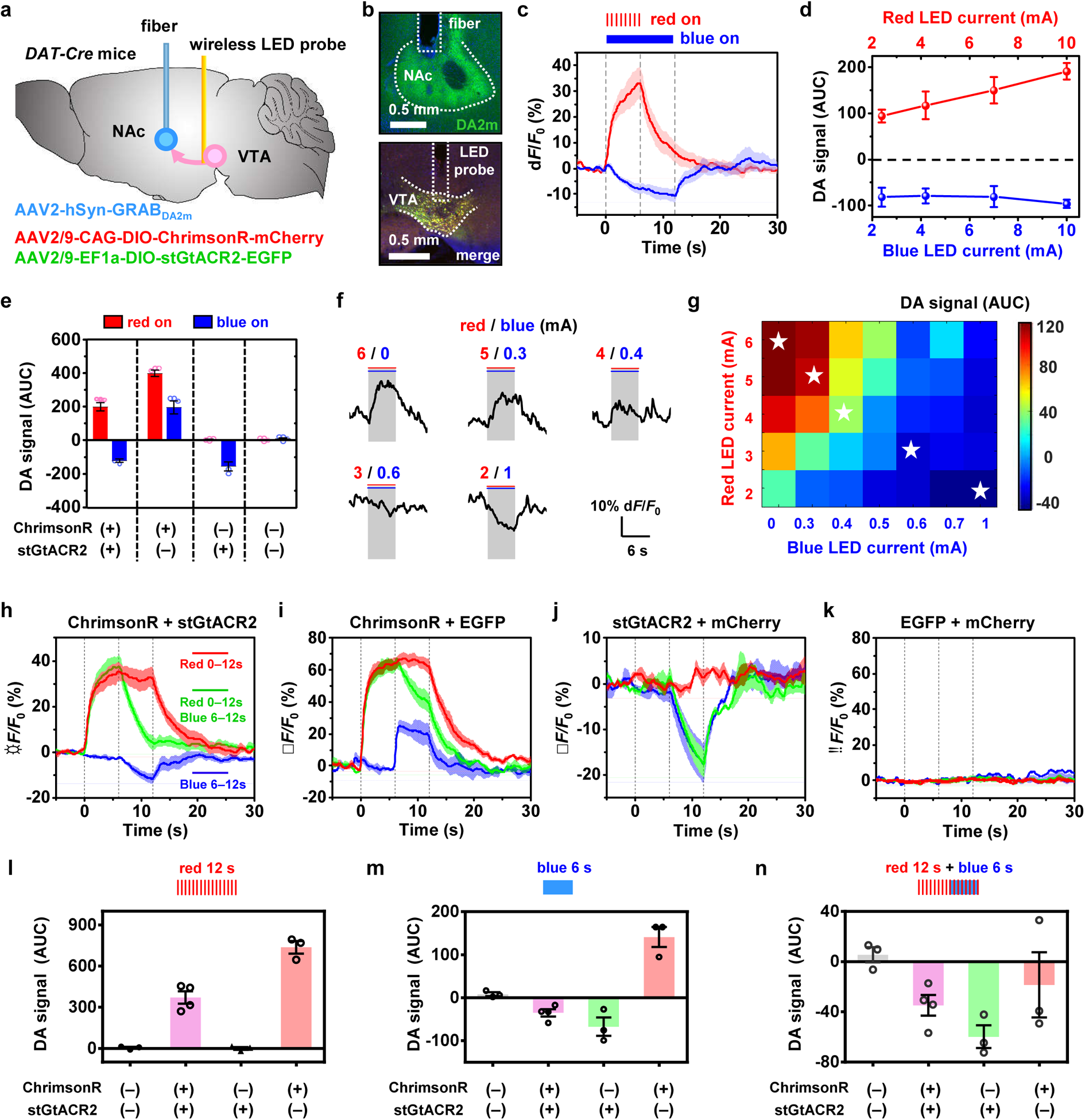
Bidirectional, *in vivo* optogenetic control of dopamine (DA) release simultaneously recorded in the nucleus accumbens (NAc) during dual-color stimulation in the VTA. **a**, Diagram illustrating the combined viral injection (AAV2/9-CAG-DIO-ChrimsonR-mCherry and AAV2/9-EF1a-DIO-stGtACR2-EGFP) followed by implanting the micro-LED probe into the VTA, and the viral injection (AAV2-hSyn-GRAB_DA2m_) followed by optical fiber implantation into the NAc of *DAT-cre* mice. **b**, Micrographs of (top) the NAc showing expression of DA2m and the fiber track, (bottom) the VTA showing co-expression of stGtACR2 and ChrimsonR (merged color) and the micro-LED probe track, 2 weeks after viral injection and probe implantation. **c**, Representative traces of DA signals in response to stimulations by the red LED (7 mA, 20 Hz, 20-ms pulse, 6 s) and the blue LED (2.4 mA, continuous, 12 s). The solid lines and shaded areas indicate the mean and s.e.m., respectively (*n* = 4 mice). **d**, Accumulative DA signals in response to red and blue stimulations under different LED currents. AUC: area under the curve during stimulation. **e**, Summary of measured DA signals (AUC) in response to red or blue stimulations (Red LED: 20 Hz, 20-ms pulse, 10 mA, 6 s; Blue LED, continuous, 3 mA, 12 s) for different mouse groups expressing ChrimsonR + stGtACR2 (n = 4 mice), ChrimsonR + EGFP (n = 3 mice), stGtACR2 + mCherry (n = 3 mice), EGFP + mCherry (n = 3 mice). Student’s t test. **f**, Example traces of DA signals under combined red and blue stimulations with varied LED currents. **g**, Summarized plot of measured DA signals (AUC) (mean value) as a function of red and blue LED currents for 6 s (*n* = 3 mice). The stars indicate results for five traces shown in Fig. 5f. **h–k**, Plots of optogenetically evoked DA transients in response to different stimulation patterns (Red: red LED on, 20 Hz, 20-ms pulse, 10 mA, 0–12 s; Blue: blue LED on, continuous, 3 mA, 6–12 s; Green: red LED on 0–12 s and blue LED on 6–12 s), for different mouse groups expressing different opsins. **h**, ChrimsonR + stGtACR2 (*n* = 4 mice). **i**, ChrimsonR + EGFP (*n* = 3 mice). **j**, stGtACR2 + mCherry (*n* = 3 mice). **k**, EGFP + mCherry (*n* = 3 mice). The solid lines and shaded areas indicate the mean and s.e.m., respectively. **l–n**, Summary of measured DA signals (AUC) for different mouse groups. **l**, red LED on, 20 Hz, 20-ms pulse, 10 mA, 0–12 s. **m**, blue LED on, continuous, 3 mA, 6–12 s. **n**, red LED on 0–12 s and blue LED on 6–12 s.

Figure 5c plots temporally resolved dopamine release in response to stimulations caused by red and blue micro-LEDs in the same mice. Other representative curves are presented in Figure S22, with results summarized in Figure 5d. Illuminating the VTA by red or blue micro-LEDs (with injection currents from 0 to 10 mA) clearly induces increased or decreased dopamine signals in the NAc, which correlates with neuron responses at the cellular level (Figure 2). Experiments operated among mouse groups expressing different opsins are also performed, with results compared in Figure 5e. In contrast to the group expressing both opsins (ChrimsonR and stGtACR2), mice with only ChrimsonR show increased dopamine release under both red and blue illuminations, and dopamine suppression is observed under blue illumination for the mice with only stGtACR2. The control group (with EGFP and mCherry) exhibits no response under optical stimulations. These results evidently explain *in vivo* preference and avoidance behaviors presented in Figure 3, showcasing the capability for precise control of neural activities based on bidirectional, dual-color modulations.

We further investigate the level of the released dopamine with coincident red and blue LED radiations. Figure 5f plots example traces of dopamine transients, in response to the red and blue micro-LEDs under pulse and continuous operations (6-s duration) with varied currents. Statistical data for accumulative dopamine concentration variations (unit: area under the curve during stimulation, or AUC) are summarized in Figure 5g and Figure S23. Decreasing the power of the red LED and increasing the power of the blue LED to cause the decline of dopamine release. Additionally, the blue LED current required to suppress the dopamine release is much lower than that of the red LED required for dopamine level rises, which further conforms with the cellular activities reported in Figure 2. It should be noted that the *in vivo* results in Figure 5g are not quantitatively identical to the *in vitro* electrophysiological ones in Figure 2m. In fact, higher irradiances for blue LEDs are needed for effective inhibition *in vivo*. This may be due to the fact that blue light has higher absorption in tissue and stronger illumination is required to affect enough cells in the VTA.

Figure 5h (and more examples in Figure S24) presents dopamine dynamics in response to alternated or combined red and blue LED stimulations, for mice expressing ChrimsonR and stGtACR2. The red illumination triggers the elevated dopamine release, while the use of blue light can effectively and promptly suppress the signal. Similar illumination patterns are implemented on mice expressing different opsins (Figures 5i–5k) and results are summarized and compared in Figures 5l–5n. Evidently, red or blue illuminations bidirectionally regulate dopamine signals in the experiment group (with both ChrimsonR and stGtACR2). On the other hand, mice expressing only ChrimsonR can be excited by both red and blue light, and those with only stGtACR2 can only be inhibited by blue light and show no reaction under red light. Finally, the control group (with EGFP and mCherry) does not respond to any optical stimulation patterns. Taken together, these results clearly demonstrate that wireless dual-color LED combined with optogenetics enable efficient, bidirectional control of neuronal activity in the VTA and DA release of the terminal in the NAc. It is also noted that fluorescence signals experience a drop when the blue LED is operating during red stimulation for the mice only expressing ChrimsonR (Figure 5i). Such a response is unexpected considering the fact that blue light also activates ChrimsonR. One possible explanation is that when both pulsed red light and continuous blue light are imposed simultaneously, their activation effects on ChrimsonR are not additive but competitive. On the other hand, the application of both LEDs possibly induces additional heating effects, complicating the optical and thermal responses of ChrimsonR. Nevertheless, understanding spectral and temporal properties for these opsins requires further explorations in the future work.

## DISCUSSION

Optogenetic modulations offer a viable solution for cell-type-specific targeting, but still demand biological and engineering tools for versatile functionalities. In this work, we report that bidirectional optogenetic activation and inhibition in behaving mice can be achieved by co-expressing spectrally distinct excitatory and inhibitory channelrhodopsins, cooperating with a wireless, dual-color micro-LED probe for colocalized red and blue stimulations. With collective results at the cellular, circuit, behavioral and social levels, successful proof-of-concept demonstrations in the VTA-NAc circuitry in freely moving mice clearly establish the utility of such a synergistic methodology.

The unique characteristics of thin-film, microscale optoelectronic devices, and their integration strategies implicate prominent advantages over conventional waveguide-coupled or single-color LED-based sources. In terms of technology developments, more directions can be explored and some preliminary efforts are presented below: 1) Multiple red and blue micro-LEDs can be assembled on the implantable probe via the stacking approach, realizing multi-channel, bidirectional modulations imposed on multiple brain regions (Movie S4). It is noted that the efficacy of preference behavior regulations demonstrated in Figure 3 would also be enhanced by implementing dual-channel stimulations bilaterally in the VTA^58–61^. 2) The use of miniaturized wireless control circuits allows independently addressing multiple, untethered animal objects, making the interrogations of more complex social behaviors feasible^62–65^. 3) Micro-LEDs with other colors can also be included and target specific opsins more efficiently. For example, a tri-color probe can be achieved by similarly stacking red, green, and blue micro-LEDs in a vertical structure (Movie S5). Independently controlled micro-LEDs can interact with different opsins that express in the same or different populations of cells. 4) Micro-LED based optogenetics can also be combined with previously reported electrical, optical and chemical sensors as well as microchannel devices for drug administrations^66–69^, achieving closed-loop neural activity stimulations and detections. While arrays of blue InGaN micro-LEDs can be directly grown on silicon and integrated with recording electrodes^41^, the combination of simultaneous dual-color stimulations and electrophysiological recordings in the deep brain region like VTA require more sophisticated device design and fabrication and have not been attempted here. Considering the system dimensions, wireless circuits can be further miniaturized by employing inductive coupling-based power transfer strategies^24, 26^. In terms of biocompatibility, tissue lesions can be further reduced by using thinner substrates with much lower stiffness^45^. Taken together, we envision a multi-spectral, multi-channel, multi-objective, and multi-functional optical neural interface, realizing wireless, real-time, bi-directional, close-loop, and simultaneous interrogations of neural signals in living animals with desirable system stability and biocompatibility.

On the other hand, the efficacy of optogenetic modulations can be further optimized by advanced biological engineering. Other combinations of opsins, like ChR2 and Jaws^36^, have also been proved to be feasible. A recent effort fuses excitatory and inhibitory channelrhodopsins (ChrimsonR and GtACR2) in a single, trafficking-optimized tandem protein (BiPOLES), improving the efficiency of opsin co-expression^70^. Moreover, the combination of super sensitive opsins (Jaws^15^, ChRmine^71^, ReaChR^72^, SOUL^13^, etc.) could be explored for bidirectional modulations via extra-cranial illumination without implants. Since relatively large irradiance (> 200 mW/mm^2^) is required to penetrate into the deep tissue, corresponding light sources and control systems have to be miniaturized for wireless operation in mice. In addition to the bidirectional modulation of the same cell population, the dual-color (or even tri-color) micro-LED probe can also selectively activate genetically distinct and spatially intermingled cells expressing spectrally resolved opsins (for example, blue for ChR2 or GtACR2, green for C1V1 or GtACR1 and red for ChrimsonR or halorhodopsin)^3, 7, 8, 21^ and provide prolonged activation of neuronal activity by regulating step-function opsins (for example, SSFO, SOUL)^11, 13, 73^, which is also of critical importance for understanding the circuit structures and functions. It should be also noted that these dual-color or multi-color modulations should be carefully designed, and their efficacies are highly dependent on the spectral overlap among opsins, the irradiance of different colors, levels of viral expressions, etc. Possible applications would include bidirectional modulations of neural activities by leveraging electrophysiological or neurotransmitter signals for medications of seizures or Parkinson’s disease^29, 74^. Besides the brain system, the spinal cord and peripheral neural circuits can also be targeted for medical treatments^75–77^. In summary, the results presented here provide a viable means for fundamental neuroscience studies and advanced neuroengineering applications.

## Methods

### Device Fabrication

#### Fabrication of red and blue micro-LEDs

Details about the thin-film micro-LED structures and fabrication processes can be found in our previous work^37, 39^. Via metal-organic chemical vapor deposition (MOCVD), red and blue LEDs are originally grown on gallium arsenide (GaAs) and sapphire substrates, respectively. The red LED active device contains an indium gallium phosphide (InGaP) based multiple quantum well structure (thickness ~5.6 μm) grown on GaAs with an Al_0.95_Ga_0.05_As sacrificial interlayer. The blue LED active device contains an indium gallium nitride (InGaN) based multiple quantum well structure (thickness ~7 μm) grown on sapphire. Photolithographic process, wet/dry etching and metallic electrode deposition define the LED geometry (size ~180 μm × 125 μm). Freestanding, thin-film red LEDs are formed by selectively removing the Al_0.95_Ga_0.05_As sacrificial layer in a hydrofluoric acid (HF) based solution. Freestanding, thin-film blue LEDs are formed by a laser lift-off (with a KrF excimer laser at 248 nm). Poly(dimethylsiloxane) (PDMS) based stamps are employed to pick up released thin-film micro-LEDs and transfer them onto other foreign substrates.

#### Fabrication of thin-film filters

Details about the thin-film filter fabrication can be found in our previous work^40^. The wavelength selective thin-film filters is based on a multilayered titanium dioxide (TiO_2_) and silicon dioxide (SiO_2_) structure (thickness ~ 6.6 μm) comprising (from top to bottom): 19 nm TiO_2_ / 15 periods (89 nm SiO_2_ + 52 nm TiO_2_) / 63 nm SiO_2_ / 66 nm TiO_2_ / 22 periods of (73 nm SiO_2_ + 42.5 nm TiO_2_) / 154 nm SiO_2_. The structure is deposited on GaAs by ion beam-assisted sputter deposition. The filter shape (size ~ 180 μm × 125 μm) is defined by laser milling (Nd: YVO_4_ laser, 1064 nm). After removing the GaAs substrate in solutions (NH_4_OH: H_2_O_2_: H_2_O = 1:1:2), freestanding thin-film filters can be picked up and transferred printed onto other foreign substrates by PDMS stamps.

#### Fabrication of dual-color micro-LED probes

The process flow to fabricate the dual-color micro-LED probe is described in Figure S1. The micro-LED probe is based on a flexible substrate made of composite polyimide (PI) and copper (Cu) films (from top to bottom: Cu/PI/Cu = 18/25/18 μm, from DuPont). From bottom to top, a red LED, an optical filter and a blue LED are sequentially transferred printed on the film by PDMS in a vertical stack, via a customized alignment setup (with a lateral alignment accuracy of ~2 μm). Between each device layer, spin-coated SU-8 based epoxy (thickness ~ 2–5 μm) serves an optically transparent and electrically insulating bonding layer. Each individual LED is metalized with sputtered metal layers (Cr/Au/Cu/Au = 10/100/500/100 nm). Laser milling (ultraviolet laser 365 nm) defines the needle shape (width ~320 μm, length ~5 mm). A bilayer of PDMS (~20 μm, by dip-coating) and parylene (~15 μm, by CVD) is used for waterproof encapsulation. For comparison in Figure 1g, laterally arranged red-blue micro-LED probes are also fabricated with similar procedures.

### Device Characterization and Modeling

#### Structural and optoelectronic characterizations

Scanning electron microscopic (SEM) images are taken with a ZEISS Auriga SEM / FIB Crossbeam System. Optical images are captured by a microscope MC-D800U(C). The LED emission spectra are measured with a spectrometer (HR2000+, Ocean Optics). Current-voltage characteristics are recorded by a Keithley 2400 source meter. LED irradiances and external quantum efficiencies are measured using a spectroradiometer with an integrating sphere (LabSphere Inc.). The angular-dependent emission profiles are measured by a standard Si photodetector (DET36A, Thorlabs) with the device mounted on a goniometer.

The probe’s mechanical properties are measured with dynamic mechanical analysis (DMA Q800, TA Instruments). For *in vitro* tests, micro-LED probes are implanted into the brain phantom. The brain phantom is made of agarose (0.5% w/v), hemoglobin (0.2% w/v), and intralipid (1% w/v) mixed in phosphate buffer solutions (98.3% w/v). After stirring, the mixed liquid is heated to boiling and then naturally cooled to room temperature, forming the gels.

#### Optical modeling

A ray-tracing method based on Monte Carlo simulations (TracePro free trial version) is used to simulate the light propagation in the brain tissue. In the model, the tissue (size 5 × 5 × 5 mm^3^) has absorption coefficients of 0.2 /mm and 0.08 /mm, and scattering coefficients of 47 /mm and 35 /mm, for blue (475 nm) and red (630 nm) wavelengths, respectively. The tissue has an anisotropy factor of 0.85 and a refractive index of 1.36^78^. Planar sources similar to the surface of micro-LEDs (size 125 × 180 μm^2^) emit 10^6^ rays for each monochromic wavelength (475 nm and 630 nm), assuming a Lambertian distribution. For the stacked micro-LEDs, the blue and red sources are overlaid. For the lateral case, the two sources are adjacent to each other.

#### Thermal characterizations

For *in vitro* experiments, the micro-LED probes are implanted into the brain phantom (~0.5 mm underneath the surface, at a depth of ~5 mm), and various currents are injected into the LEDs. The temperature distributions on the phantom surface are mapped with an infrared camera (FOTRIC 228). For comparison, the temperature inside the phantom is also measured by inserting an ultra-fine flexible micro thermocouple (Physitemp, IT-24P) (~0.3 mm above the LED probe tip). For *in vivo* experiments, the micro-LED probe is bonded with the same thermocouple and inserted into the anesthetized mouse brain. Steady-state temperature rises are recorded from the thermocouple with a thermometer (Physitemp, BAT-12).

#### Thermal modeling

3D steady-state heat transfer models are established by finite element analysis (Comsol Multiphysics). Materials and corresponding parameters (density, thermal conductivity and heat capacity) used in the model include: brain tissue (1.1 g/cm^3^, 0.5 W/m/K, 3.7 J/g/K), parylene (1.2 g/cm^3^, 0.082 W/m/K, 0.71 J/g/K), PDMS (0.98 g/cm^3^, 0.16 W/m/K, 1.5 J/g/K), polyimide (1.4 g/cm^3^, 0.15 W/m/K, 1.1 J/g/K), GaN (6.1 g/cm^3^, 130 W/m/K, 0.49 J/g/K), Cu (9.0 g/cm^3^, 400 W/m/K, 0.39 J/g/K), SU-8 (1.2 g/cm^3^, 0.2 W/m/K, 1.2 J/g/K), GaP (4.4 g/cm^3^, 110 W/m/K, 0.43 J/g/K). The boundary condition is natural heat convection to air at 20 °C. The micro-LED probe serves as the heat source, with the input thermal power estimated by *P* = *V* × *I* × (1 − EQE), where *V*, *I,* and EQE are the measured voltage, current, and corresponding external quantum efficiencies for LEDs. The thermocouple is also included in the model for comparison, which is 0.3 mm above the LED probe tip. It consists of two metal wires: copper (diameter 31 μm) and constantan (45Ni-55Cu alloy, thermal conductivity 21 W/m/K, diameter 31 μm), which are encapsulated by polyester to form a wire with an outer diameter of 127 μm.

### Wireless Circuit Design

A customized circuit module is designed to wirelessly operate the implantable micro-LED probe by independently controlling the red and the blue micro-LEDs for bi-directional modulation. The schematic diagram is depicted in Figure S5, with a detailed circuit diagram, layouts, and components presented in Figures S6–S9. The core components include: a micro-controller (nRF24LE1, Nordic Semiconductor) operated at 2.4 GHz for data communication, and two LED drivers (ZLED7012, Renesas Electronics Corp.) with programmable current levels, frequencies, pulse widths for providing a constant current to micro-LEDs^38^. The driver provides programmable constant current levels ranging from 1.8 mA to 20 mA and also supports arbitrary waveforms. Two red LEDs are used on the circuit board outside the brain for signal indication, without affecting the mouse behaviors^79^. Control signals are wirelessly transmitted from the transmitting module connected to a laptop computer via an antenna operating at a radio frequency of 2.4 GHz, with a communication distance up to 50 m. Different from our previous design^80^, here a polyimide-based flexible circuit board is implemented for reduced weight. The wireless circuit has a footprint of ~2.2 cm × 1.3 cm and a weight of 1.9 g (including a 0.9-g rechargeable lithium-ion battery with a capacity of 35 mAh). Used in the experimental conditions in this paper, the battery life can reach up to 2 hours. The circuit can be connected with the implanted micro-LED probes via a flexible printed circuit (FPC) connector before the animal experiment (Figure S8).

### Biological Studies

#### Animals

Animal care is in accordance with the institutional guidelines of National Institute of Biological Sciences in Beijing (NIBS), with protocols proved by Institutional Animal Care and Use Committee (IACUC). All mice are socially housed in a 12 h/12 h (lights on at 8 am) light/dark cycle, with food and water *ad alibitum*. Wildtype, male C57BL6/N mice are purchased from VitalRiver (Beijing, China). *DAT-Cre* (B6.SJLSlc6a3tm1.1(cre) Bkmn; JAX Strain 006660) and *CamkIIa-Cre* (B6.Cg-Tg(Camk2a-cre)T29-1Stl/J; JAX Strain 005359) mice are at least six weeks old at the time of surgery.

#### Virus production

The AAV2/9-CAG-DIO-ChrimsonR-tdTomato (Plasmid #130909) vector^7^ is from Addgene. The AAV2/9-EF1a-DIO-stGtACR2-EGFP-Kv2.1 plasmid is synthesized and constructed according to the original reports^8, 9^. The AAV-hSyn-GRAB_DA2m_ plasmid is provided by Prof. Yulong Li at Peking University^55^. AAV vectors are packaged into serotype2/9 vectors, which consist of AAV2 ITR genomes pseudotyped with AAV9 serotype capsid proteins. AAV vectors are replicated in HEK293 cells with the triple plasmid transfection system and purified by chloroform^81^, resulting in AAV vector titers of about 5×10^12^ particles/mL. Virus suspension is aliquoted and stored at −80 °C. AAV titers are measured using real-time qPCR. Adeno-associated viral particles are produced at the Vector Core Facility at Chinese Institute for Brain Research, Beijing (CIBR).

#### Stereotaxic Surgery

Adult *DAT-Cre* mice are anesthetized with an intraperitoneal injection of Avertin (250 mg/kg) before surgery and then placed in a standard stereotaxic instrument for surgical implantation (Figure S12). After disinfection with 0.3% hydrogen peroxide, a small incision of the scalp is created to expose the skull. Then, 0.3% hydrogen peroxide is applied to clean the skull. A small craniotomy is made, followed by carefully removing the dura with a thin needle. A calibrated pulled-glass pipette (Sutter Instrument) is lowered to the VTA (coordinates 3.64 mm posterior to the bregma, 0.5 mm lateral to the midline, and 4.6 mm ventral to the skull surface). The virus is delivered through a small durotomy by a glass micropipette using a microsyringe pump (Nanoliter 2000 injector with the Micro4 controller, WPI).

Four different combinations of virus are used for *in vivo* experiments:

1. ChrimsonR + stGtACR2 (AAV2/9-CAG-DIO-ChrimsonR-tdTomato and AAV2/9-EF1a-DIO-stGtACR2-EGFP-Kv2.1, 1:1);
2. mCherry + EGFP (AAV2/9-EF1a-DIO-mCherry and AAV2/9-EF1a-DIO-EGFP, 1:1);
3. ChrimsonR + EGFP (AAV2/9-CAG-DIO-ChrimsonR-tdTomato and AAV2/9-EF1a-DIO-EGFP, 1:1)
4. stGtACR2 + mCherry (AAV2/9-EF1a-DIO-stGtACR2-EGFP-Kv2.1 and AAV2/9-EF1a-DIO-mCherry, 1:1)

A bolus of 0.4 μL of mixed virus is injected into the VTA at 46 nL/min. The pipette is held in place for 10 min after the injection and then slowly withdrawn. For the real-time place preference (RTPP) behavioral test, the dual-color LED probe implantation is carried out after virus injection and diffusion. The probe was slowly implanted at the side of the target brain areas with an angle of 45° to midline (coordinates 3.64 mm posterior to the bregma, 0.8 mm lateral to the midline, and 4.85 mm ventral to the skull surface). The probe is fixed to the skull with dental cement. For dopamine detection with fiber photometric recording, the AAV-hSyn-GRAB_DA2m_ (0.2 μL) is injected into the NAc (coordinates: 1.18 mm front to the bregma, 0.75 mm lateral to the midline, and 3.75 mm ventral to the skull surface). Optical fiber implantation is carried out after virus injection and diffusion. A piece of optical fiber (FT200UMT, 200 μm O.D., 0.39 NA, Thorlabs) is fit into an LC-sized ceramic fiber ferrule (Shanghai Fiblaser, China) and implanted over the target brain areas with the tip 0.1 mm above the virus injection sites in the NAc. The ceramic ferrule is fixed to the skull with Cyanoacrylate adhesive (TONSAN 1454) and dental cement. After the virus injection and device implantation, mice are housed individually to prevent potential damage to the probe. All subsequent experiments are performed at least 2 weeks after injection to allow sufficient time for transgene expression and animal recovery.

#### Immunohistochemistry

The micro-LED probes are implanted into the VTA region and fixed to the skull with Cyanoacrylate adhesive (TONSAN 1454) and dental cement. After 1 day or 3 weeks following the surgery, mice are sacrificed with an overdose of pentobarbital and transcardially perfused with 0.1M phosphate buffer saline and 4% paraformaldehyde. After pcryoprotection in 30% sucrose, brains are post fixed in 4% PFA for 4 hours and 30% sucrose for 24 hours, 40-μm-thick coronal sections are prepared by a cryostat (Leica CM1950). After 5-min PBS rinses for 3 times, the sections are blocked in PBST (PBS + 0.3% Triton X-100) with 3% bovine serum albumin for 1 hour, and incubated in rabbit anti-Iba1 (1:1000, Wako, Cat.#019-19741) and chicken anti-GFAP antibody (1:2000, Sigma, Cat#AB5541) dissolved in PBST at 4 °C for 24 hours. After 5-min washes in PBS for 5 times, the sections are incubated with Alexa Cy3-conjugated goat anti-rabbit antibody (1:500, Jackson ImmunoResearch) and Alexa 488 goat anti-chicken antibody (1:500, Thermo Fisher Scientific) dissolved in PBST for 2 hours at room temperature followed by 5-min washes in PBS for 3 times. Finally, the samples are cover slipped in 50% glyceroland slides and photographed with an automated slide scanner (VS120 Virtual Slide, Olympus) or a confocal scanning microscope (DigitalEclipse A1, Nikon). The numbers of DAPI, stGtACR2 and ChrimsonR stained cells or the numbers of Iba1+ microglia and GFAP+ astrocytes are analyzed with Fiji software (https://fiji.sc/) and their percentages in cell population indicated by DAPI are calculated.

#### In vitro optogenetics evaluation

Electrophysiological properties of the neurons in brain slices are measured with whole-cell patch-clamp recording technique following a previously described procedure^31^. After two weeks of virus expression (ChrimsonR + stGtACR2), mice are deeply anesthetized with pentobarbital (100 mg/kg i.p.) and intracardially perfused with 5 mL ice-cold oxygenated modified artificial cerebrospinal fluid (ACSF) at a rate of 2 mL/min. After perfusion with modified ACSF containing (in mM): 225 sucrose, 119 NaCl, 2.5 KCl, 0.1 CaCl_2_, 4.9 MgCl_2_, 1.0 NaH_2_PO_4_, 26.2 NaHCO_3_, 1.25 glucose, 3 kynurenic acid, and 1 Na L-ascorbate (Sigma-Aldrich), brains are quickly removed and placed in ice-cold oxygenated ACSF containing (in mM): 110 choline chloride, 2.5 KCl, 0.5 CaCl_2_, 7 MgCl_2_, 1.3 NaH_2_PO_4_, 25 NaHCO_3_, 20 glucose, 1.3 Na ascorbate, and 0.6 Na pyruvate. Sagittal sections (200 μm thick) are prepared using a Leica VT1200S vibratome. Brain slices are incubated for 1 hour at 34 °C with ACSF saturated with 95% O_2_/5% CO_2_ and containing (in mM) 125 NaCl, 2.5 KCl, 2 CaCl_2_, 1.3 MgCl_2_, 1.3 NaH_2_PO_4_, 25 NaHCO_3_, 10 glucose, 1.3 Na ascorbate, and 0.6 Na pyruvate. The internal solution within whole-cell recording pipettes (3–6 MΩ) contains (in mM): 130 K-gluconate, 10 HEPES, 0.6 EGTA, 5 KCl, 3 Na_2_ATP, 0.3 Na_3_GTP, 4 MgCl_2_, and 10 Na_2_-phosphocreatine (pH 7.2–7.4). Voltage-clamp recordings are performed using a MultiClamp 700B amplifier (Molecular Devices). Traces are low-pass filtered at 2.6 kHz and digitized at 10 kHz (DigiData 1440, Molecular Devices). The data are acquired and analyzed using Clampfit 10.0 software (Molecular Devices). Red and blue illuminations are provided by a fiber-coupled to two laser sources (633 nm and 473 nm).

#### In vivo extracellular electrophysiology recordings

We perform the extracellular single-unit recording as our previously described procedure^82^. Adult CamkIIa-Cre mice are anesthetized with an intraperitoneal injection of Avertin (250 mg/Kg). A bolus of 1 μL of virus (AAV2/9-CAG-DIO-ChrimsonR-tdTomato and AAV2/9-EF1a-DIO-stGtACR2-EGFP-Kv2.1, 1:1) is injected into neocortex and hippocampus at different depths (AP −1.6 mm, ML 1.1 mm, DV 0.4 to 2.0 mm increments; 200 nL/site). A silver wire (127 μm diameter, A-M system) is attached to a skull-penetrating M1 screw above the olfactory bulb to serve as the ground reference. For head-fixed preparations, a custom-made titanium head-plate is secured to the skull with dental acrylic. Recordings are performed 2 weeks after viral injection. Mice are held in place with a header bar cemented to the skull. The dual-color micro-LED probe is lowered into the neocortex with a 15° angle from medial to lateral and another 10° angle from anterior to posterior. 8 stereotrodes with Pt-Ir wires (diameter 17.5 μm, platinum 10% iridium, California Fine Wire, USA) are used for extracellular electrophysiology recordings. The distance between the stereotrodes and the micro-LEDs is ~500 μm. To reduce the noise induced by the stray capacitance coupling from LEDs to electrodes, we use a tinfoil shield to encompass the electric circuit of the LED. Voltage signals are digitized and recorded by an Open-Ephys board (http://www.open-ephys.org/). Signals from each recording electrode are band-pass-filtered between 300 and 6000 Hz and sampled at 25 kHz. Single units are discriminated with principal component analysis (Offline sorter, spike2, CED, UK). Recorded units smaller than 2.5 times the noise band or in which interspike intervals are shorter than 2 ms are excluded from further analysis. Red light (pulse width 0.2 s, interval 3 s, 50–100 pulses, LED current 10 mA) and blue light (pulse width 1 s, interval 3 s, 50–100 pulses, LED current 1 mA) are used to activate or inhibit neurons. Data are analyzed and plotted with a custom-developed MATLAB program.

#### Calcium imaging in acute brain slice

Mice are injected with 0.4 μL of Raav2/9-hsyn-GCaMP6m into the S1. After 2 weeks of recovery, acute brain slices are prepared. Ca^2+^ fluorescent signals in GCaMP6m expressing cells are captured with a 20× water immersion objective on a confocal microscope (FV1000, Olympus). For cell activation, 1 μM α-amino-3-hydroxy-5-methyl-4-isoxazolepropionic acid (AMPA, No.A6816, Sigma-Aldrich) is added in ACSF to enhance the Ca^2+^ signals. Control experiments are performed by applying pure ACSF. Fluorescence images are analyzed via ImageJ. Normalized fluorescence changes are calculated as d*F/F = (F − F*_0_*) / F*_0_, where *F*_0_ is the baseline intensity (before 10 s).

#### Real-time place preference test

After 2 weeks of recovery and AAV expression in *DAT-cre* mice, real-time place preference/avoidance (RTPP) tests are performed in a custom-made two-compartment apparatus. The RTPP apparatus comprises two white plastic chambers (size: 25 × 25 × 30, L × W × H in cm), separated by half-open opaque plastic walls in the middle (8 × 30, L × H cm). An RTPP test consists of 3 phases: 10-min baseline (pretest), 30-min red LED stimulation, 30-min rest (no stimulation), and 30-min blue LED simulation. During the pretest phase, individual mice are allowed to freely explore the entire apparatus for 10 min. Mice that exhibit a strong initial preference or avoidance for one of the side chambers (preference index > 60% or < 40%) are excluded from further experiments. During the red LED stimulation phase, mice that entered the left chamber received red light pulses (20 Hz, 20-ms pulse, 7 mA). During the blue light stimulation phase, mice that entered the left chamber receive constant blue light (continuous, 5 mA). The locomotion of mice is assessed from the video recording data using a custom-developed MATLAB program.

#### Social interaction test

After 2 weeks of recovery and AAV expression, social interaction tests are performed with *DAT-cre* mice. Two or three mice are placed in an open-field arena (30 × 30 cm^2^), wireless dual-color micro-LED probes provide red (20 Hz, 20-ms pulse, 7 mA) and blue (continuous, 5 mA) stimulations, and their behaviors are recorded by a camera. In paired-mice experiments, the two mice receive the same 3-phase stimulations: 1) 5-min baseline (pretest, no stimulation), 5-min red LED stimulation, 5-min free interaction (no stimulation), and 5-min red LED simulation; 2) 2-hour rest in individual housing; 3) 5-min baseline (pretest, no stimulation), 5-min blue LED stimulation, 5-min free interaction (no stimulation), and 5-min blue LED simulation. For triple-mice interactions, the three mice receive the following stimulations: 5-min baseline (pretest, no stimulation), 5-min LED stimulation (red, red, blue), 5-min free interaction (no stimulation), and 5-min LED simulation (blue, blue, red). The tests are repeated three times, by alternating the order. Experiment groups express stGtACR2 + ChrimsonR, and control groups express EGFP + mCherry. The videos are analyzed by Etho Vision XT 15 and social behaviors including sniff and attack are manually labeled and scored.

#### Fiber photometry for dopamine (DA) detection

Details for the fiber photometric system are described previously^82^. The excitation source is a 488 nm semiconductor laser (Coherent, Inc. OBIS 488 LS, tunable power up to 60 mW). A dichroic mirror with a 452–490 nm reflection band and a 505–800 nm transmission band (Thorlabs, Inc. MD498) is employed for wavelength selection. A multimode fiber (Thorlabs, Inc., 200 μm in diameter and 0.39 in numerical aperture) coupled to an objective lens (Olympus, 10×, NA 0.3) is used for optical transmission. The fluorescence signals are collected with a photomultiplier tube (PMT) (Hamamatsu, Inc. R3896) after filtering by a GFP bandpass filter (Thorlabs, MF525-39). An amplifier (C7319, Hamamatsu) is used to convert the current output from the PMT to voltage signals, which are further filtered through a lowpass filter (40 Hz cut-off; Brownlee 440, USA). The fluorescence signals are digitalized at 100 Hz and recorded by a Power 1401 digitizer and Spike2 software (CED, UK). Photometric data are exported and analyzed with MATLAB. The fluorescence signals (Δ*F*/*F*_0_) are processed by averaging the baseline signal *F*_0_ over a 4.5-s long control time window and presented as heatmaps or per-event plots. AUC (area under the curve) of the DA signal is used for statistical analysis. AUC is the integral of a curve that describes the variation of DA signals during light stimulation.

## Acknowledgements

This work is supported by National Natural Science Foundation of China (NSFC) (61874064), Beijing Municipal Natural Science Foundation (4202032), Tsinghua University Initiative Scientific Research Program, Center for Flexible Electronics Technology at Tsinghua University, Beijing National Research Center for Information Science and Technology (BNR2019ZS01005), CAS Key Laboratory of Brain Connectome and Manipulation (Lab No.2019DP173024, Project No.KFKT2020006). We thank Prof. Y. Li (Peking University) for support on GRAB_DA2m_, Y. Zhou and Y. Ju (Chinese Institute for Brain Research, Beijing) for support on stGtACR2.

## Author contributions

X. S. and M. L. developed the concepts. L. Li, G. T., Y. Z., X. C., Z. S., H. D., C. L., Y. X., H. W., L. Y. and X. S. performed material, device and circuit design, fabrication and characterization. L. Li, X. C., D. C, and X. S. performed the simulations. L. Li, L. Lu, Y. R., G. T., X. F., L. Y., M. L., and X. S. designed and performed biological experiments. L. Li, L. Lu, M. L. and X. S. wrote the paper in consultation with other authors.

## Data and materials availability

All data needed to evaluate the conclusions in the paper are present in the paper and/or the Supplementary information.

---

**Figure S1.**
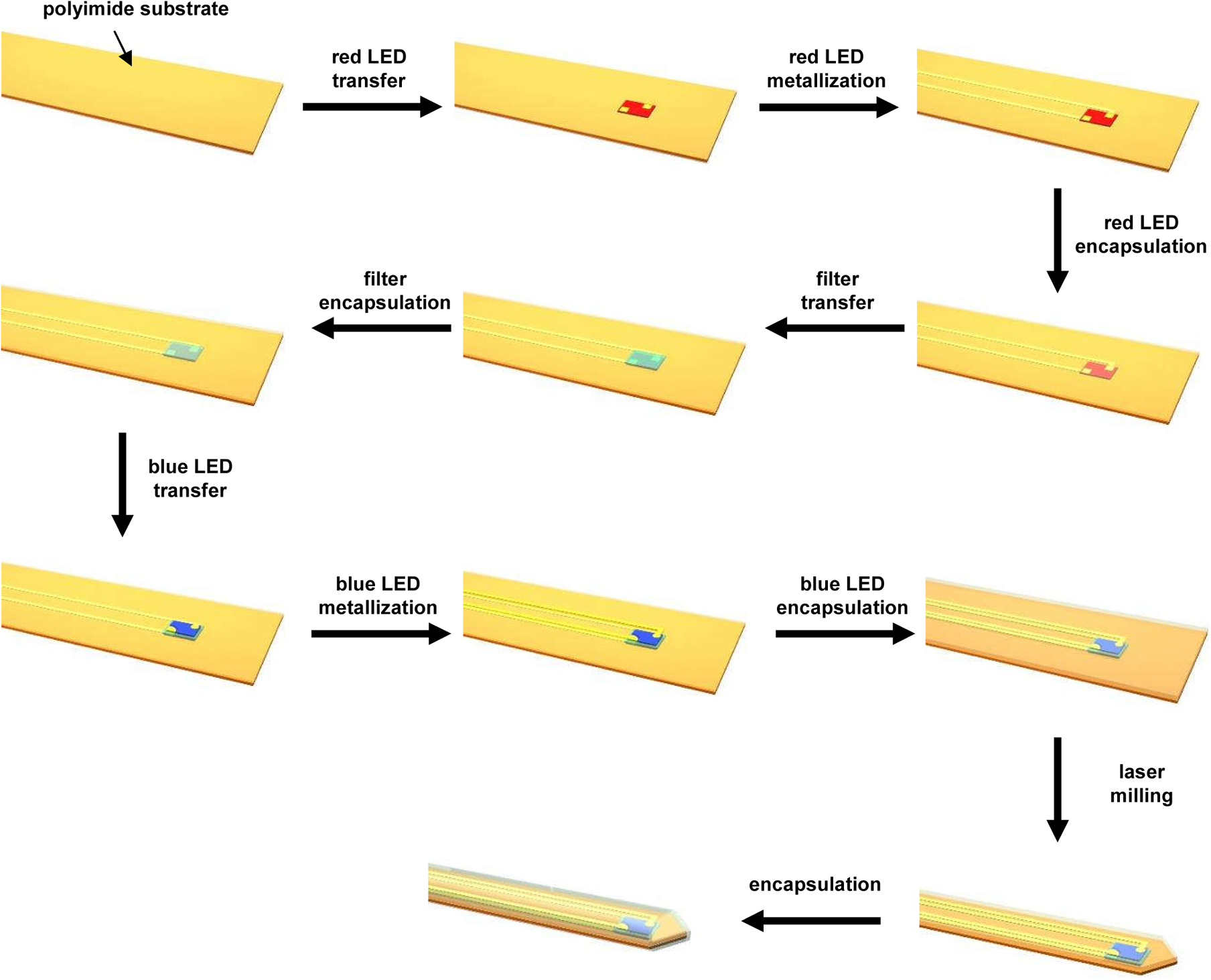
Schematic illustration of the process flow for fabricating the dual-color micro-LED probes, with vertically stacked red LED, filter and blue LED printed on PI substrates.

**Figure S2.**
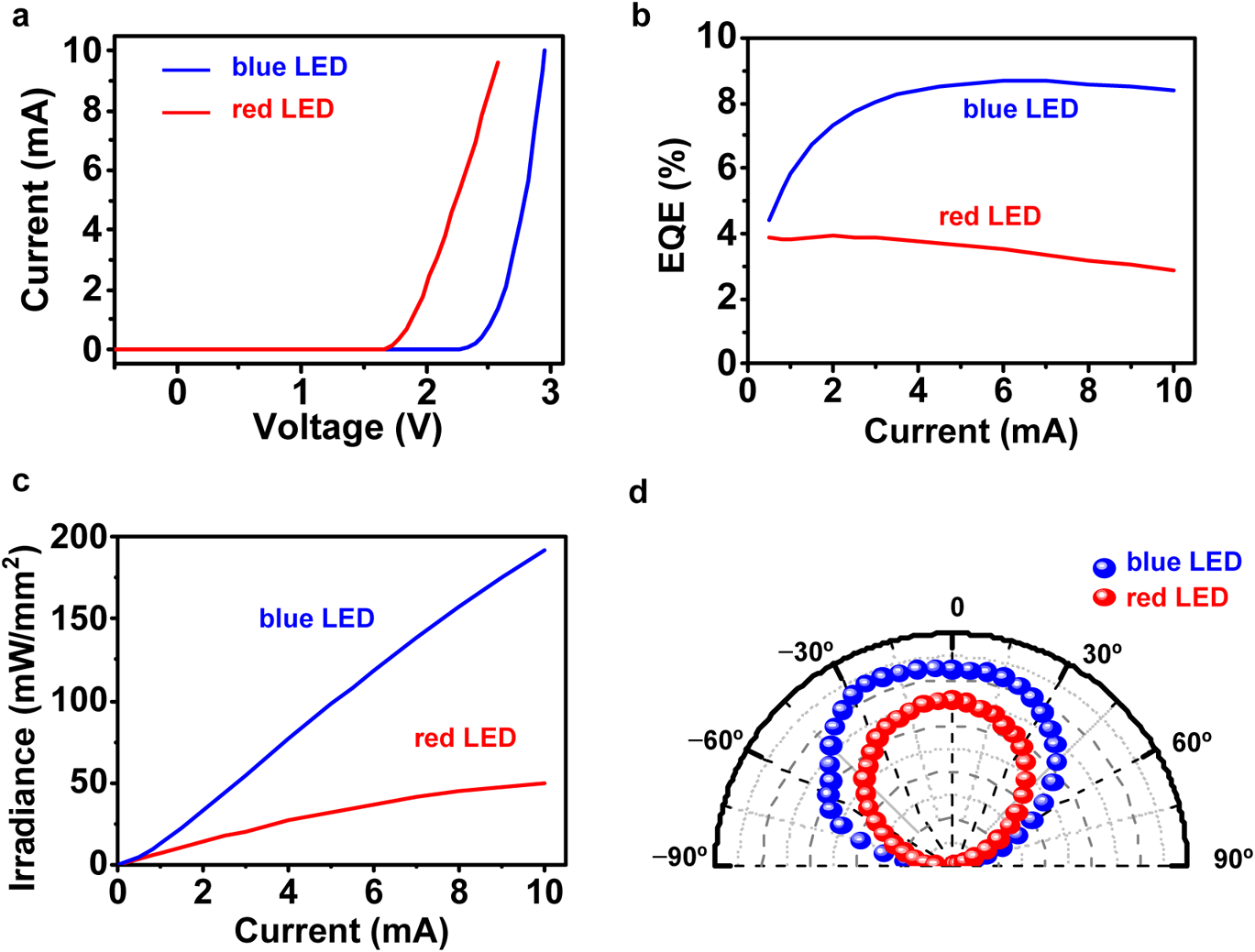
Measured optoelectronic properties of the red and the blue micro-LEDs in a probe structure. (a) Current–voltage curves. (b) External quantum efficiencies (EQEs) as a function of current. (c) Irradiance (power density on the LED surface) versus current. (d) Angular dependent emission profiles (in arbitrary unit).

**Figure S3.**
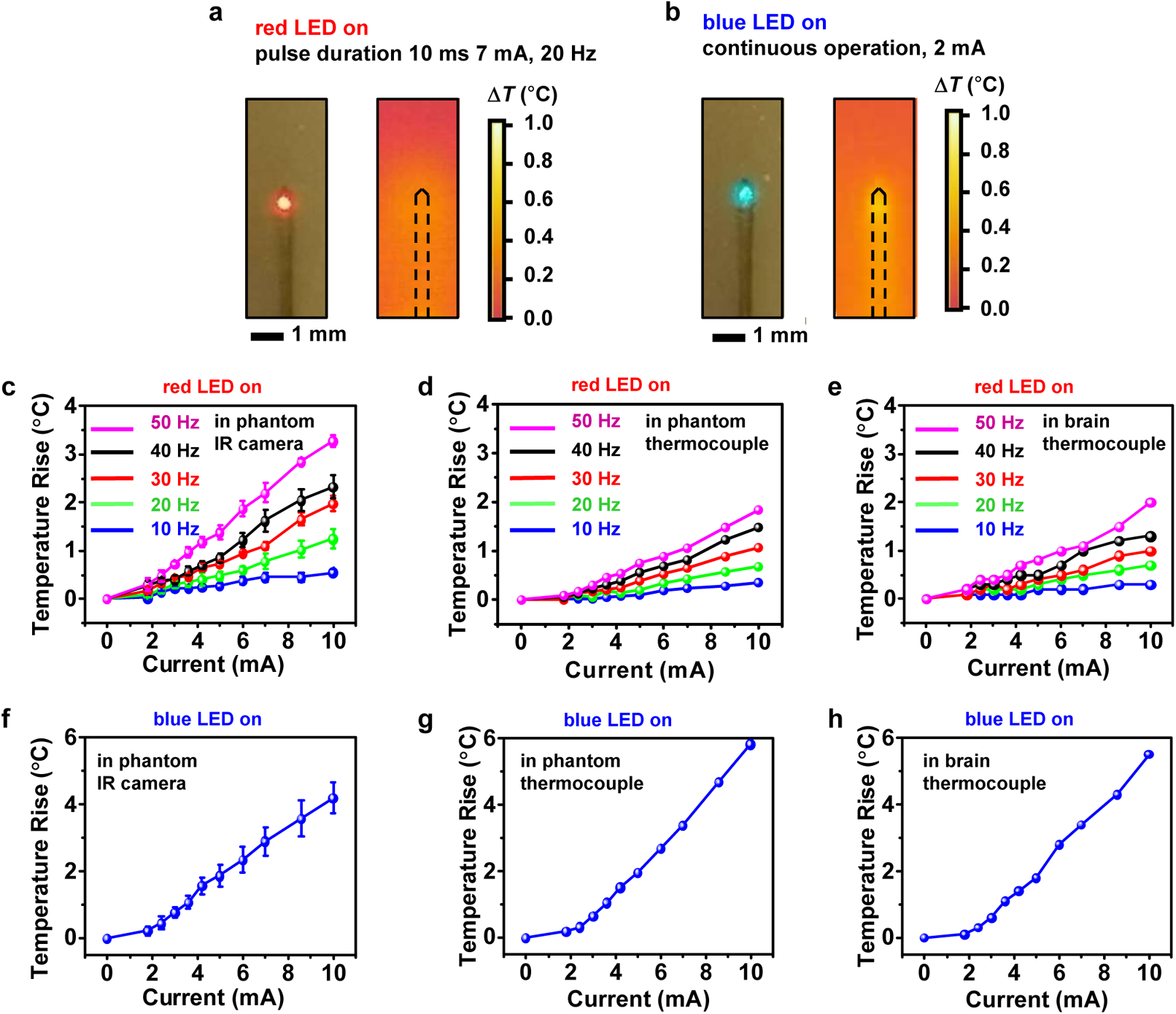
Measured thermal behaviors of a dual-color micro-LED probe. (a, b) Images and infrared (IR) thermographs showing a probe embedded into a brain phantom (~0.5 mm below the phantom surface). (a) Red LED on (injection current 7 mA, pulse frequency 20 Hz, pulse duration 10 ms). (b) Blue LED on (injection current 2 mA, continuous operation). (c) Measured temperature rise on the phantom surface by IR camera. (d) Measured temperature rise in the phantom by thermocouple. (e) Measured temperature rise in the brain of living mice by thermocouple. Red LED current 0–10 mA, frequency 10–50 Hz, pulse duration 10 ms). (f–h) Corresponding results when the blue LED is on (continuous mode).

**Figure S4.**
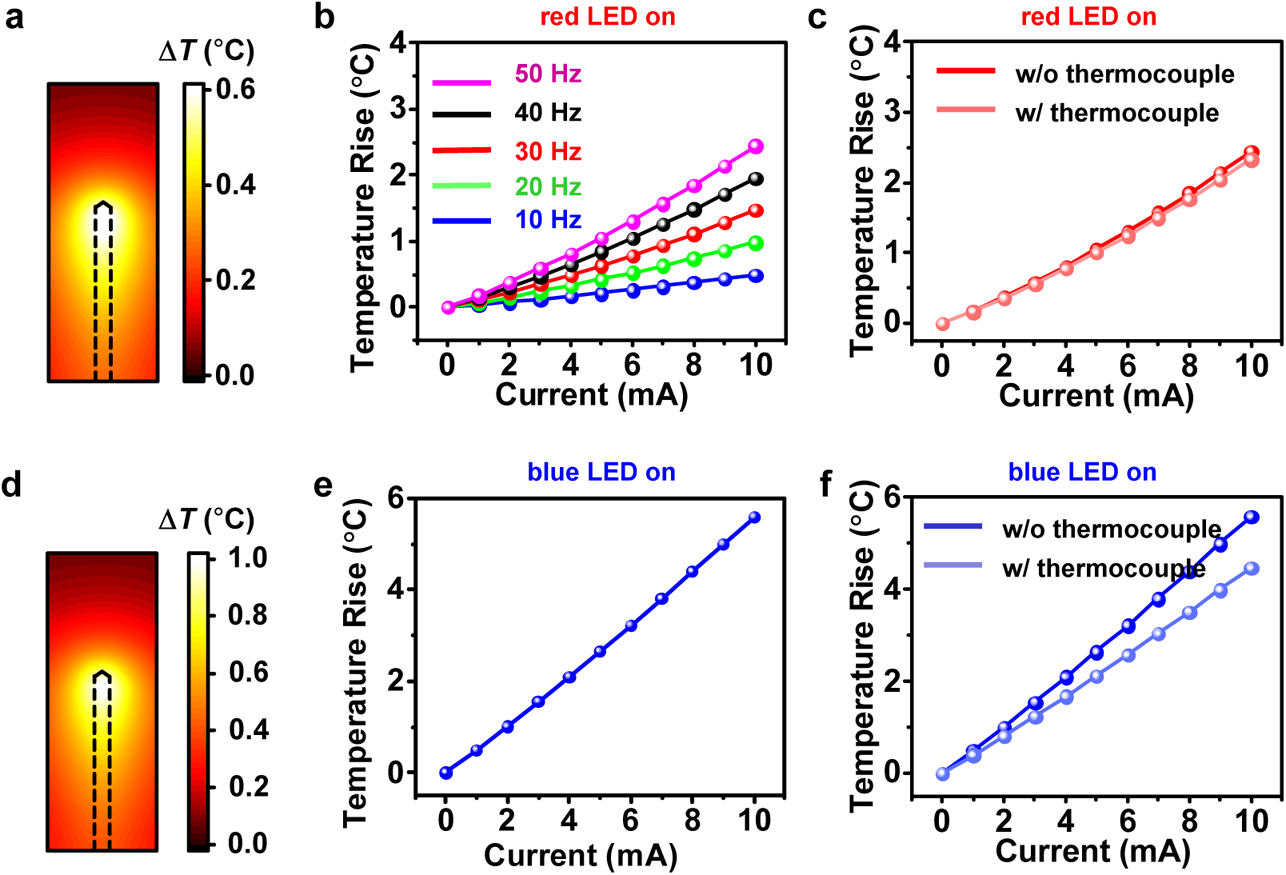
Simulated thermal behaviors of a dual-color micro-LED probe. (a) Simulated steady-state temperature distribution in the brain tissue surrounding the probe, when the red LED is on (current 7 mA, frequency 20 Hz, pulse duration 10 ms). (b) Simulated maximum temperature rise of the brain tissue when the red LED is in different operation conditions. (c) Simulated maximum temperature rise of the brain tissue when the red LED is on (frequency 50 Hz, pulse duration 10 ms). The results for models with and without the thermocouple are compared. (d–f) Corresponding results when the blue LED is on (continuous mode), current = 2 mA in (d).

**Figure S5.**
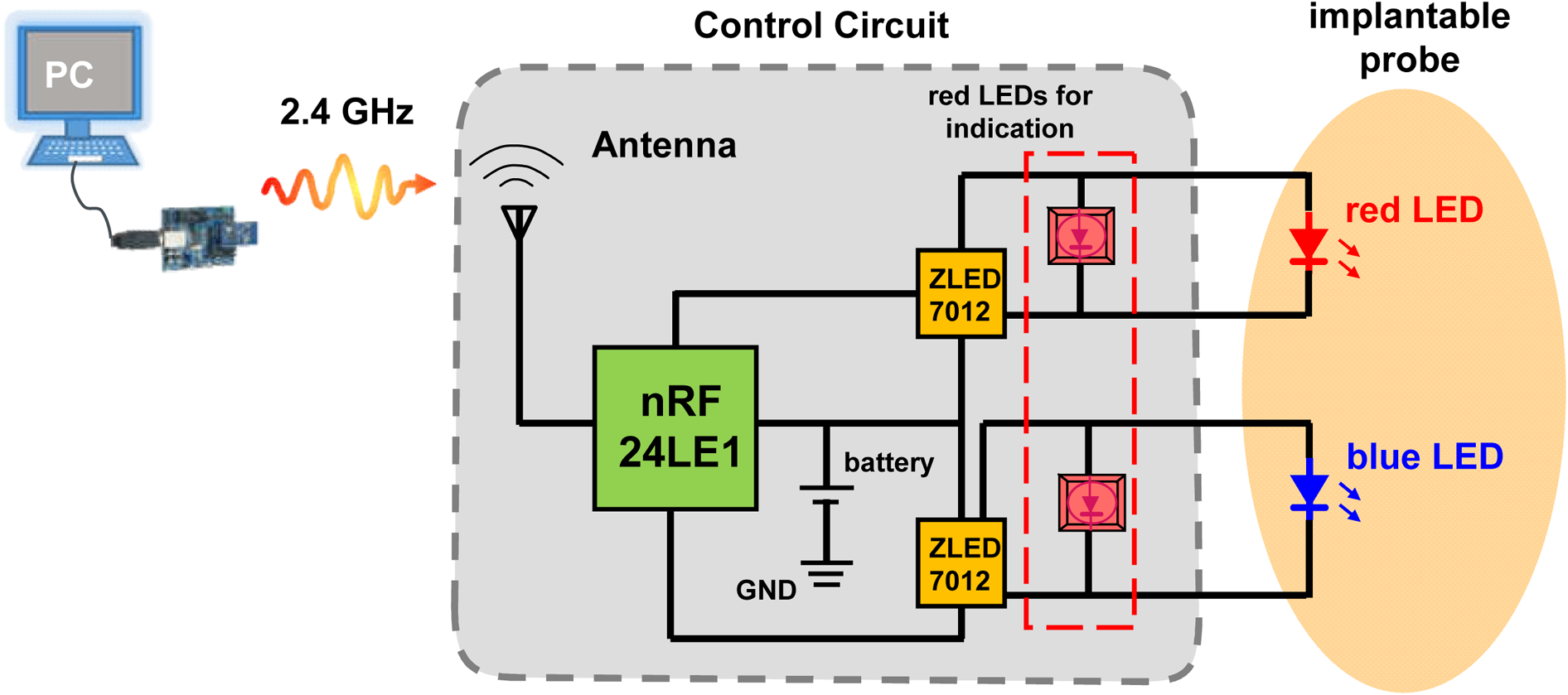
Schematic diagram showing the operational principle for the wireless circuit system controlling the dual-color micro-LED probe.

**Figure S6.**
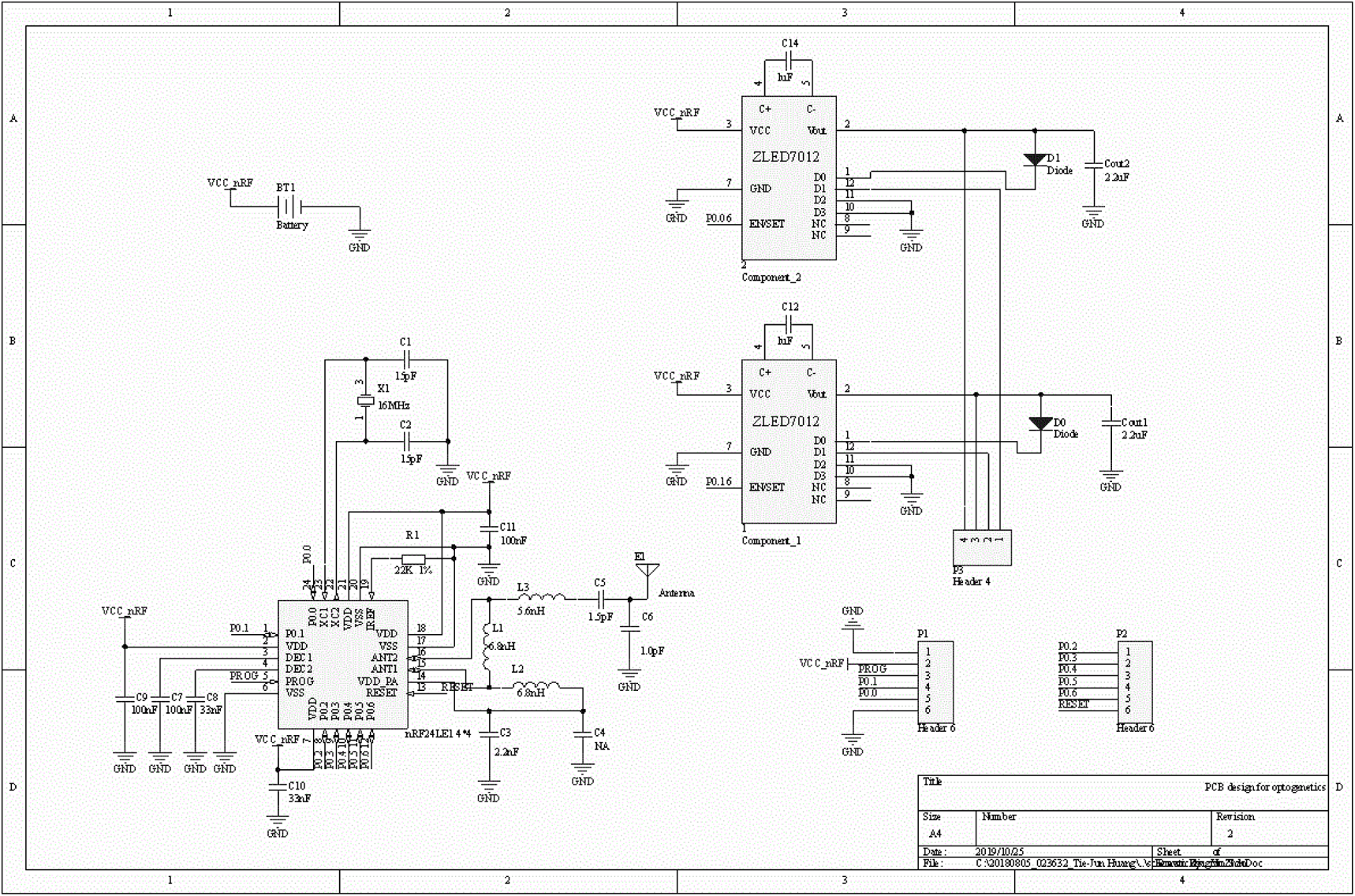
Detailed circuit design diagram of the wireless circuit system.

**Figure S7.**
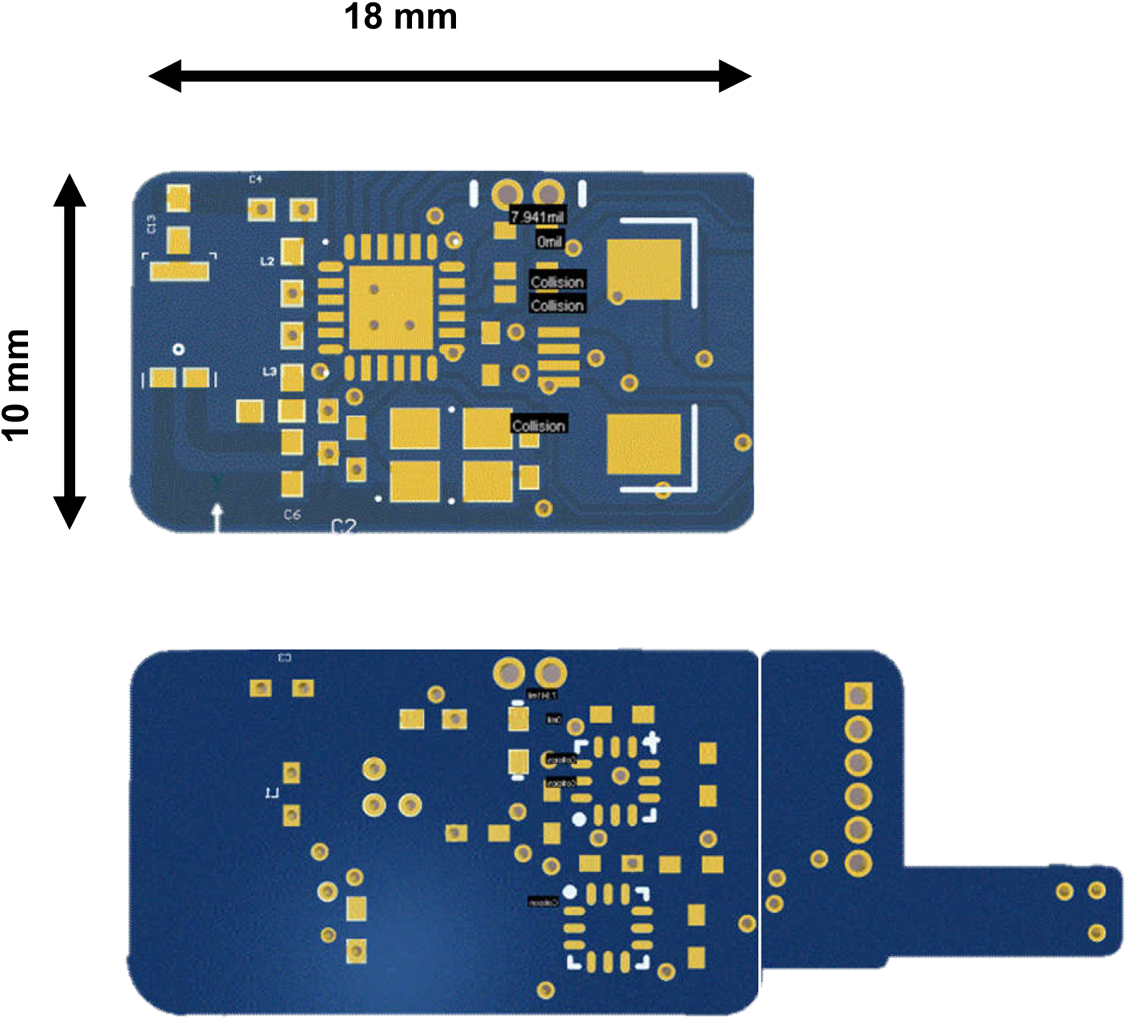
Layout of the designed printed circuit board (top: front view; bottom: back view).

**Figure S8.**
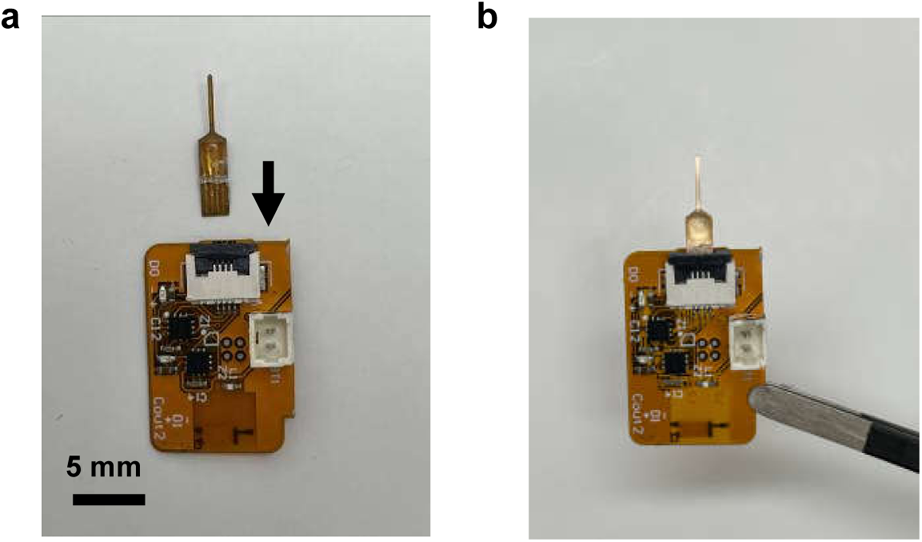
Optical images of a micro-LED probe and a control circuit. (a) before and (b) after connection via a standard 4-pin connector.

**Figure S9.**
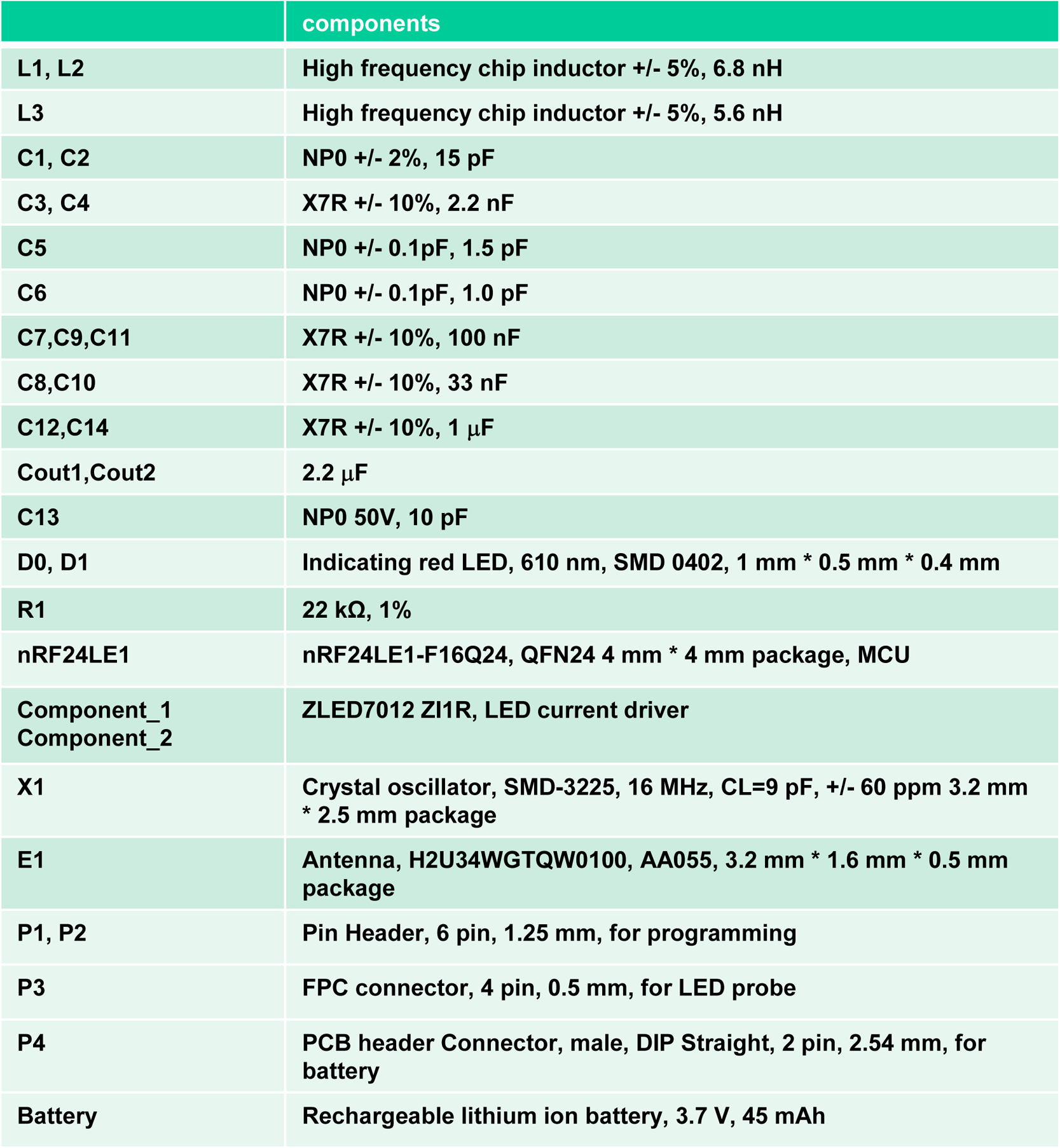
List of components used in the wireless circuit system.

**Figure S10.**
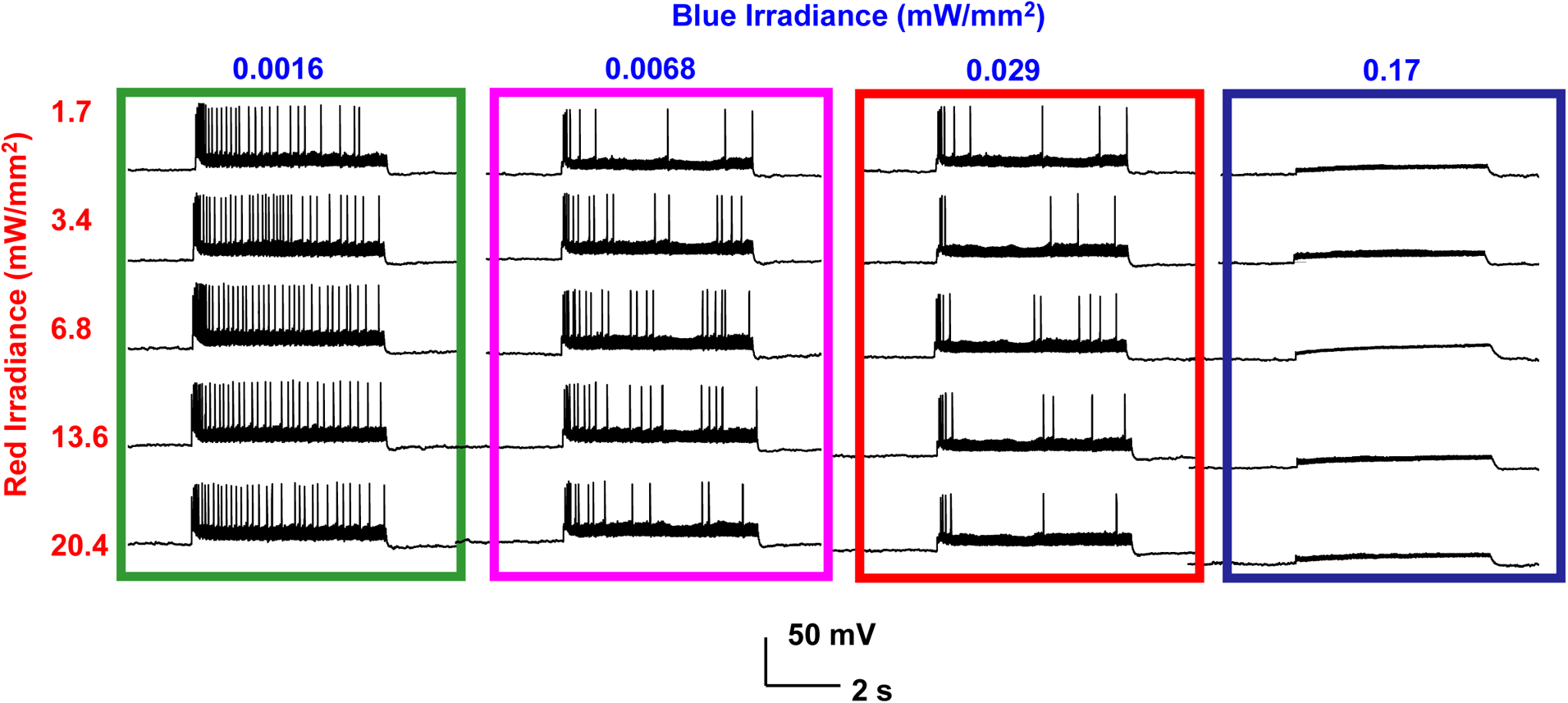
Recorded traces of a representative neuron with combined red and blue illumination at different irradiances. Red LED: 20 Hz, 10-ms pulse; Blue LED: continuous operation.

**Figure S11.**
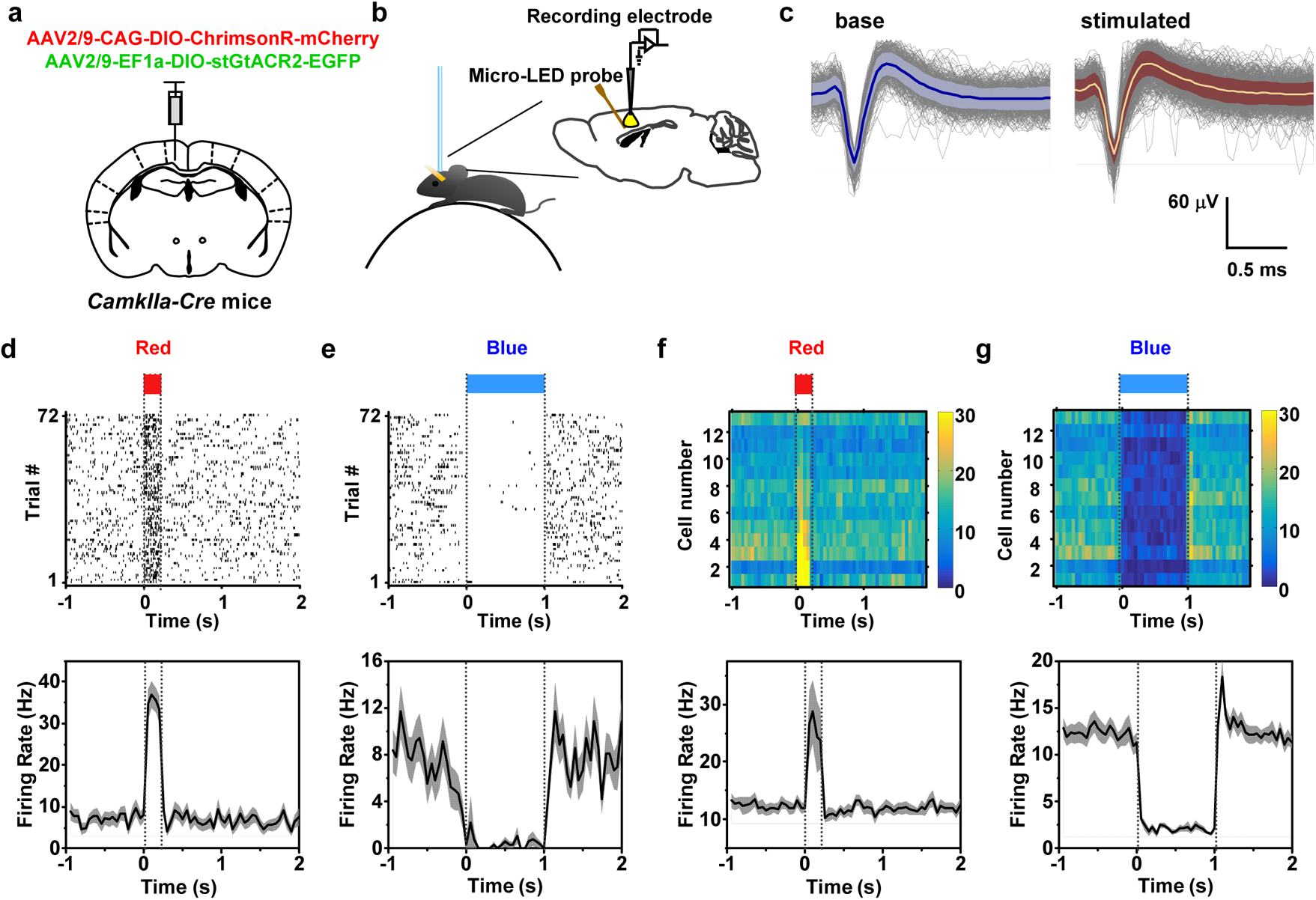
Bidirectional, *in vivo* optogenetic modulation of neural activities with dual-color illuminations in the cortex, combining with electrophysiological recordings. (a) Schematic strategy for co-expressing ChrimsonR and stGtACR2 in the primary somatosensory cortex of *CamkIIa-Cre* mice. (b) Illustration of the setup for simultaneous optogenetic stimulation and electrophysiological recordings by implanting the micro-LED probe and metal electrodes into the cortex of head-fixed mice. (c) Waveforms of a single unit recorded during baseline period before the red LED illumination (left) and during the red LED illumination period (right). Correlation coefficient between the two sets of waveforms is 0.9965. (d, e) Raster plots (top) and peri-stimulus time histogram (PSTH) plots (bottom) recorded for an example unit (*n* = 72 trials) for a sample unit during (d) red illumination (pulse width 0.2 s, LED current 10 mA) and (e) blue illumination (pulse width 1 s, LED current 1 mA). (f, g) Summarized results (top: heatmaps; bottom: PSTH plots) for multiple cells collected during (f) red illumination and (g) blue illumination (*n* = 13 cells from 2 mice, 50–100 trials for each cell).

**Figure S12.**
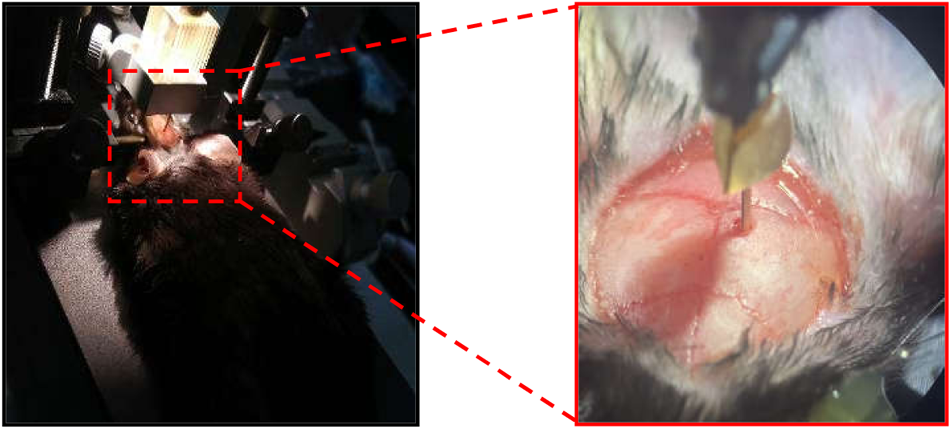
Photographs of the surgical procedure for probe implantation. A hole is created on the exposed skull of a mouse by drilling, and the dura is carefully removed by needle. The micro-LED probe is fixed on a holder controlled by a stereotaxic instrument, and slowly inserted into the targeted region.

**Figure S13.**
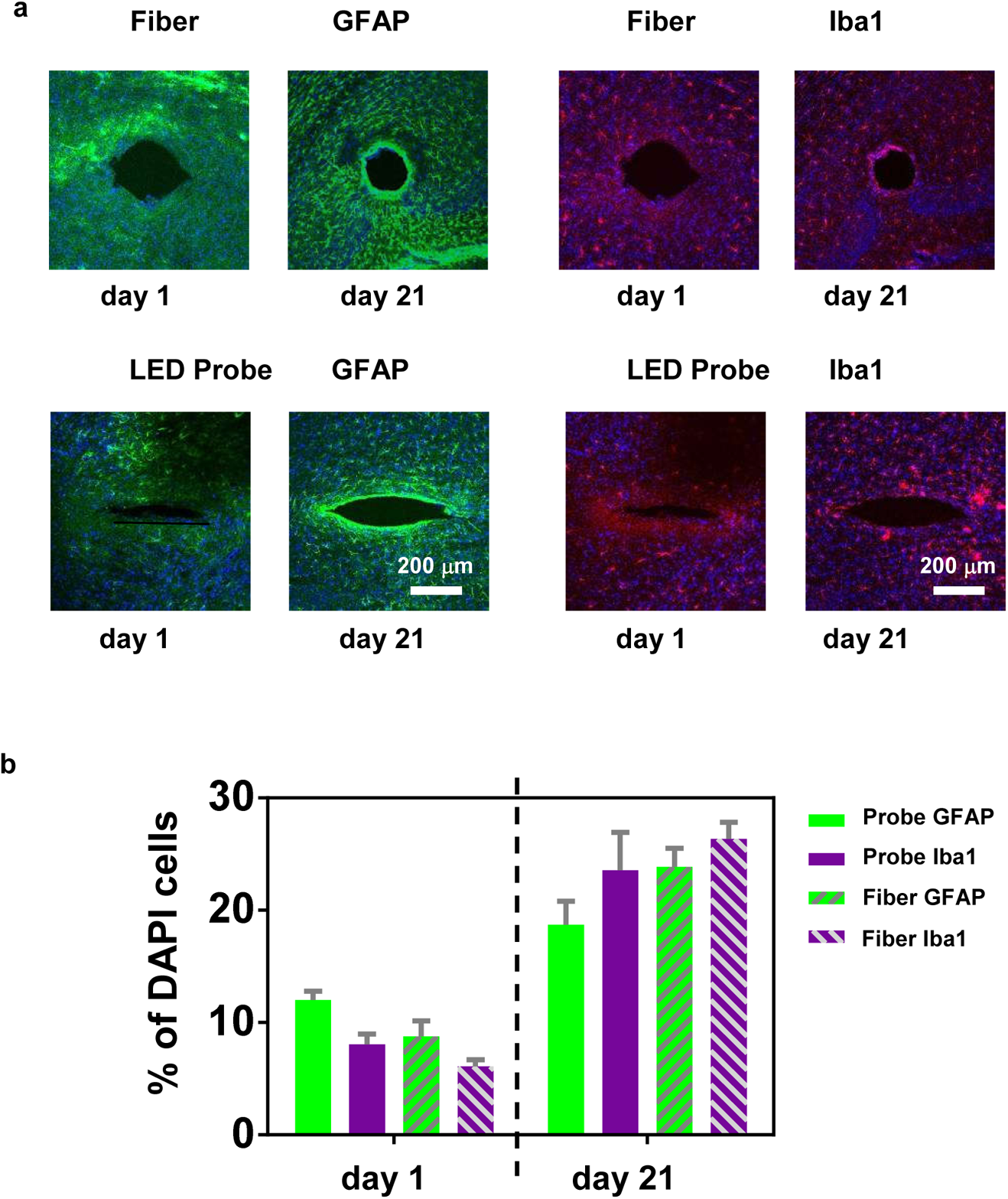
Immunoreactivity results showing the biocompatibility of the micro-LED probe, in comparison with those obtained with a silica based optical fiber. (a) Representative confocal fluorescence images of horizontal brain slices, showing immunohistochemical staining of astrocytes (GFAP) and activated microglia (Iba1) for both the LED probe and the fiber, after 1 day and 21 days implantation. Green: GFAP; Red: Iba1; Blue: DAPI. (b) Percentages of GFAP and Iba1 cell populations among DAPI cells collected at a distance of 200 μm from the edge of implantation. The LED probe and the fiber show similar results of inflammatory glial responses occurring after implantation. (*n* = 3 mice for each group).

**Figure S14.**
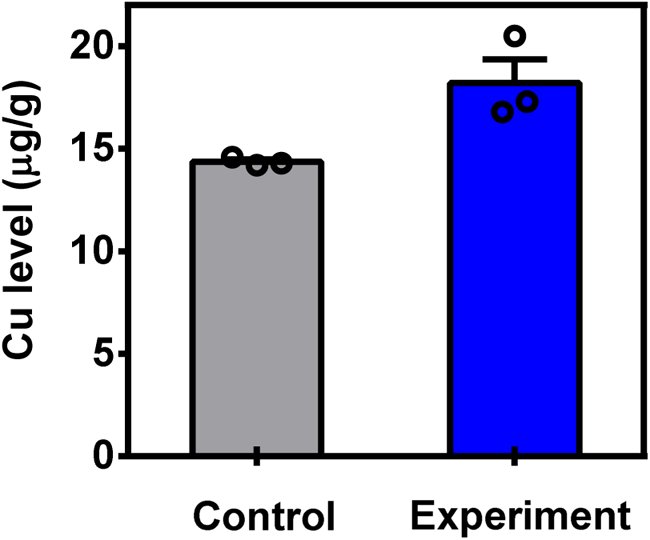
Measured Cu concentration in the brain tissue. The brain tissue is dried and analyzed with inductively coupled plasma mass spectrometry (Thermo ICP-MS iCAPQ, ThermoFisher, USA). The measured Cu level is 18.2 ± 2.3 μg/g for mice 5 weeks post probe implantation (experiment group). The result for the control group without probe is 14.4 ± 0.2 μg/g. The Cu levels in both cases are in the normal range (< 20 μg/g in mice brain, and 30– 100 μg/g in human brain). *n* = 3 mice for each group. References: Chen Y, Wang L, Geng J-H, Zhang H-F, Guo L. Apolipoprotein E deletion has no effect on copper-induced oxidative stress in the mice brain. Biosci Rep 2018, 38(5). Harrison WW, Netsky MG, Brown MD. Trace elements in human brain: Copper, zinc, iron, and magnesium. Clinica Chimica Acta 1968, 21(1): 55-60.

**Figure S15.**
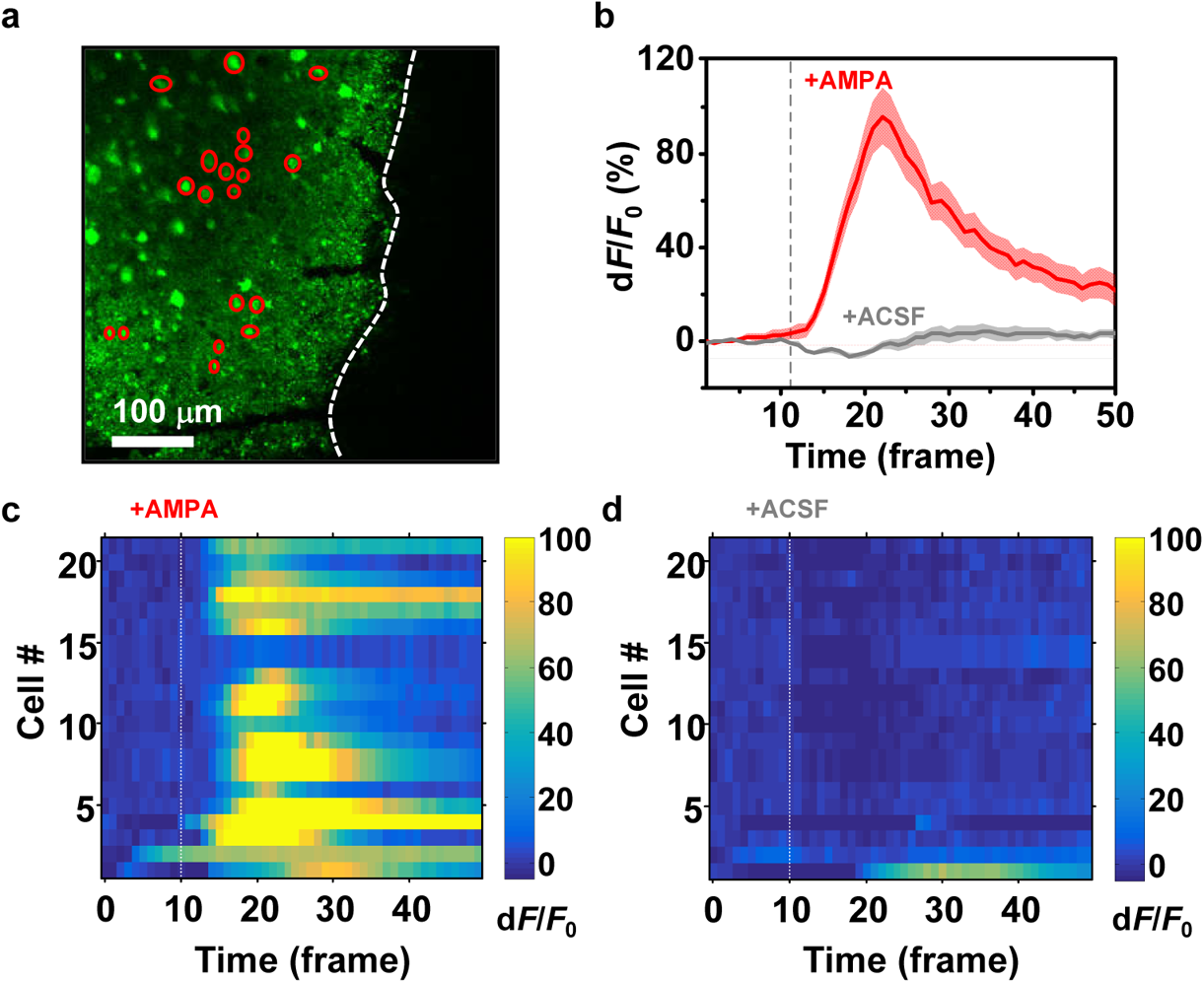
Imaging cellular calcium dynamics in acute brain slice. (a) Representative fluorescence image showing GCaMP6m-expressing cells near the probe region after 14 days implantation. Red circles mark active cells followed by AMPA administration. The dashed line marks the border of the lesion area. (b) Averaged calcium signal traces (d*F/F*_0_) for all marked neurons after applying AMPA (red line) or pure ACSF (grey line) at frame 11, the sampling rate is 1.1 s / frame. (c, d) Heatmaps showing fluorescence variations in all 21 neurons after (c) applying AMPA or (d) pure ACSF.

**Figure S16.**
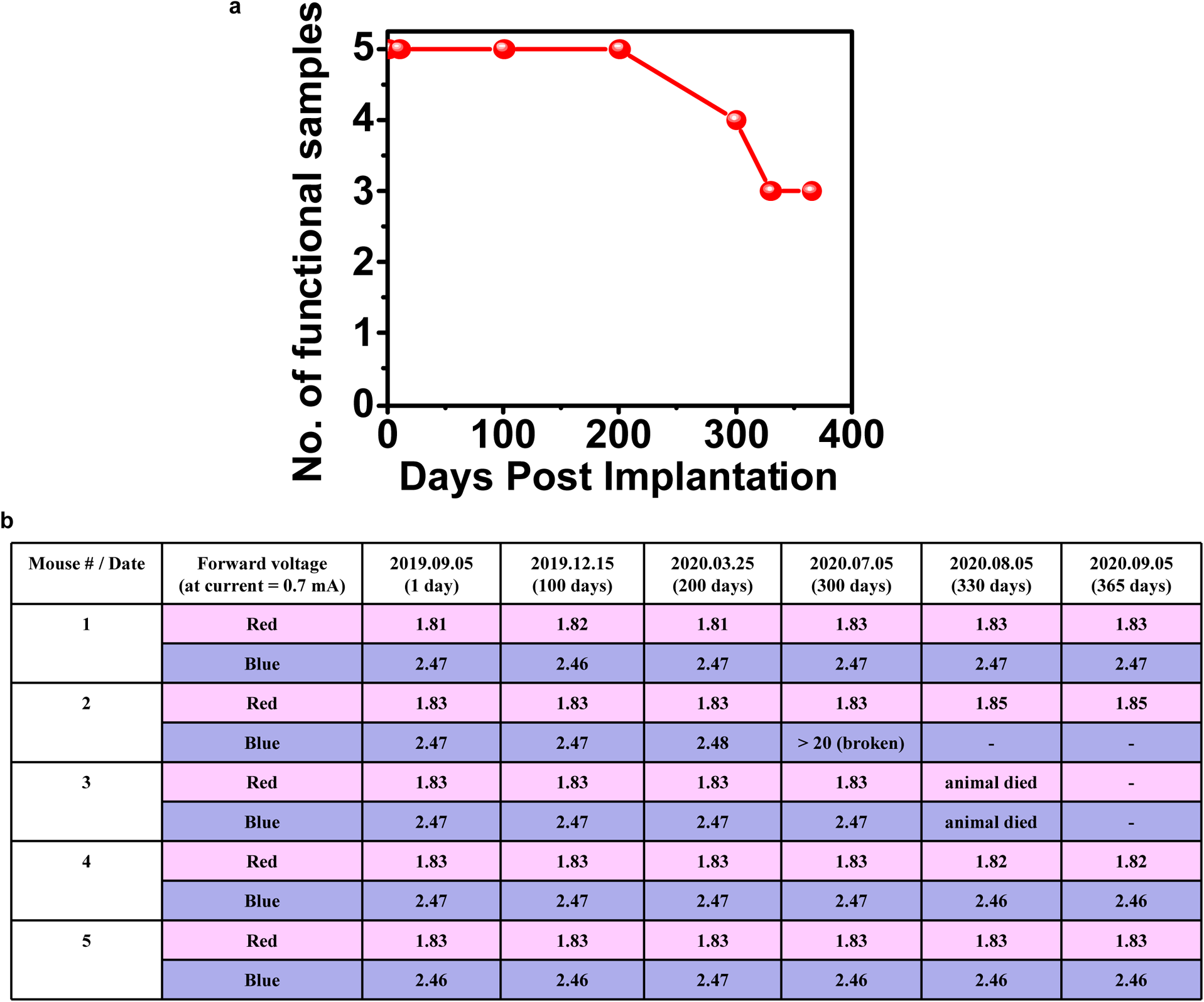
(a) Chronic stability for the dual-color micro-LED probe. 5 probes are separately implanted into 5 behaving mice. Probes with both red and blue micro-LEDs operating in the normal condition are defined as “functional probes”. (b) Summary of the performance for red and blue micro-LEDs in each probe. These probes are kept within the mouse brain. Their performance are evaluated by measuring LED’s forward voltages at a constant current of 0.7 mA. After 365 days, the functional probes are taken out and still operate normally.

**Figure S17.**
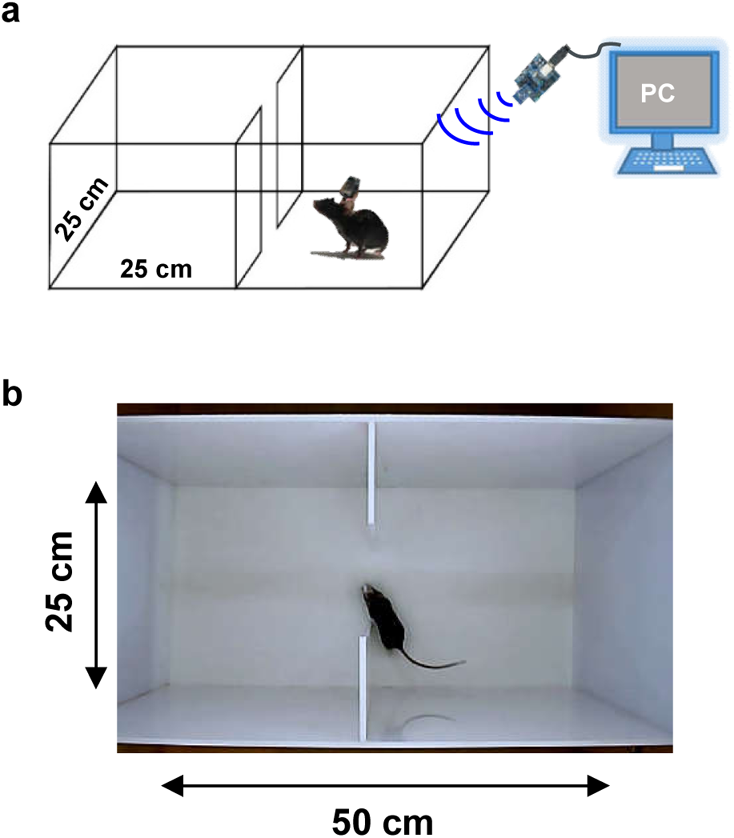
(a) Schematic diagram and (b) top-view photograph of the real-time place preference test with a two-compartment arena.

**Figure S18.**
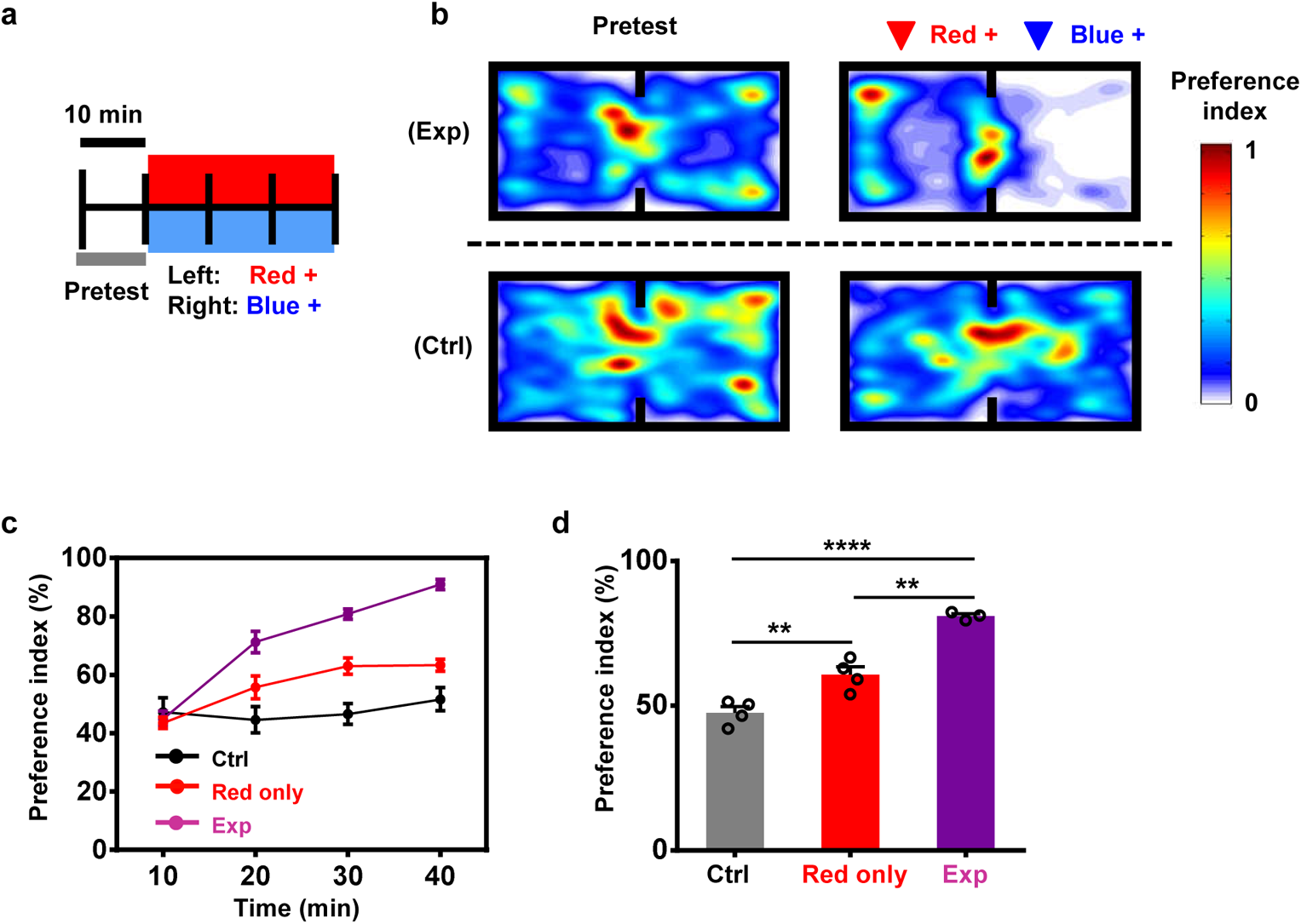
(a) Patterns used for optogenetic modulations, including a 10-min pretest, a 30-min red LED stimulation (20 Hz, 10-ms pulse, current 7 mA) in the left chamber and blue LED stimulation (continuous, current 5 mA) in the right chamber. (b) Representative heat maps comparing pretest and real-time preference behavior following both red (left chamber) and blue (right chamber) stimulation for mice expressing stGtACR2 + ChrimsonR (experiment group) and EGFP + mCherry (control group). (c) Preference indices measured at different times for mice under only red stimulations (red line, *n* = 4 mice), or red stimulations in the left chamber and blue stimulations in the right chamber (exp, purple line, *n* = 3 mice), and the control group with the red stimulation in the left chamber and blue light in the right chamber (ctrl, black line, *n* = 4 mice). (d) Summary of preference indices (the ratio of the time that mice spend in the left chamber to the whole recorded time) for mice under only red stimulations (*n* = 4 mice), or red stimulations in the left chamber and blue stimulations in the right chamber (*n* = 3 mice) for both experiment and control groups. Student’s *t* test, ** *P* < 0.01, **** *P* < 0.0001.

**Figure S19.**
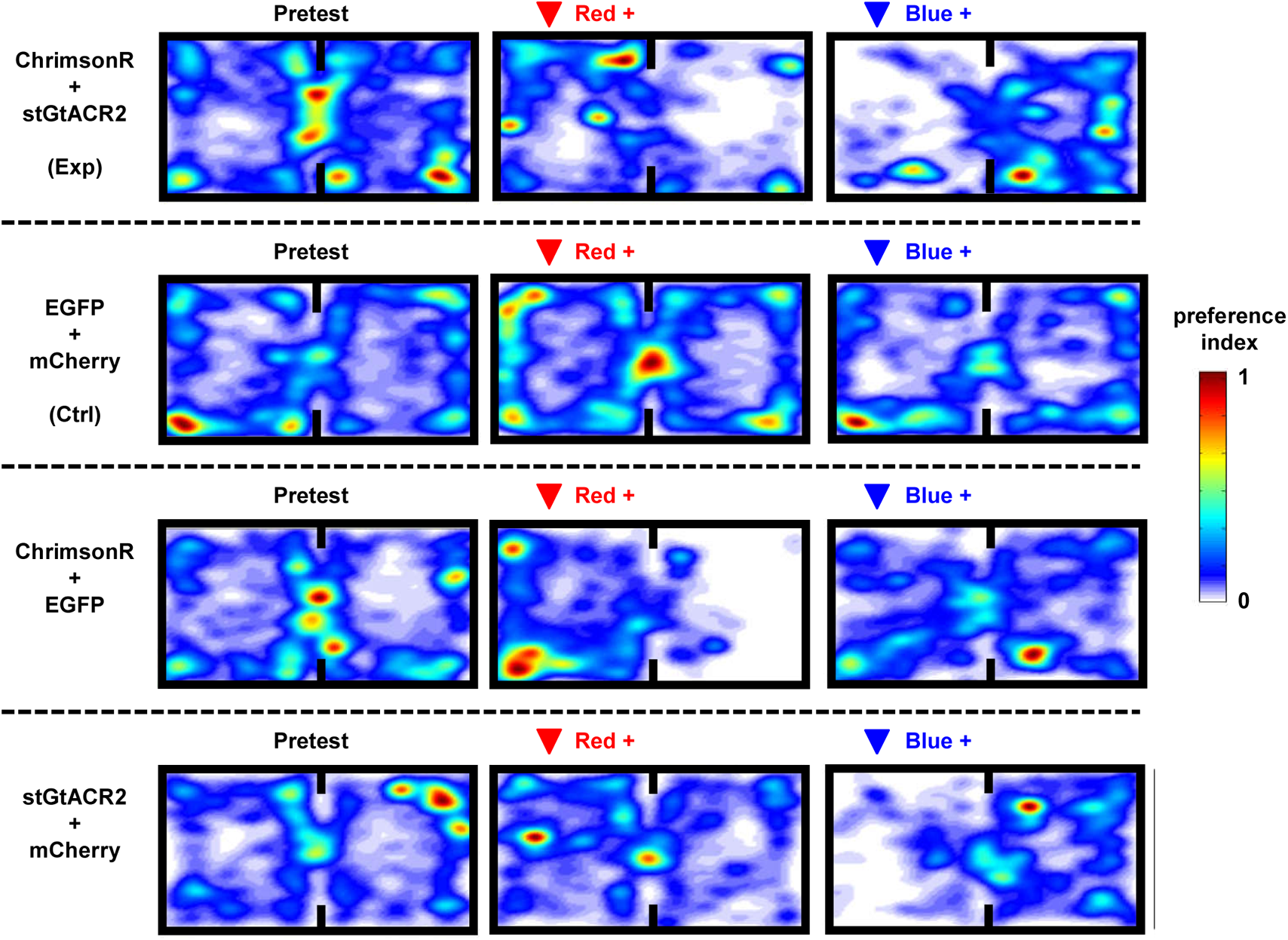
Representative heat maps showing real-time preference and aversion behavior following red or blue stimulation for mice expressing stGtACR2 + ChrimsonR (experiment, or Exp), EGFP + mCherry (control, or Ctrl), ChrimsonR + EGFP and stGtACR2 + mCherry. The data for ChrimsonR + stGtACR2 and EGFP + mCherry groups are identical to those presented in Fig. 3e.

**Figure S20.**
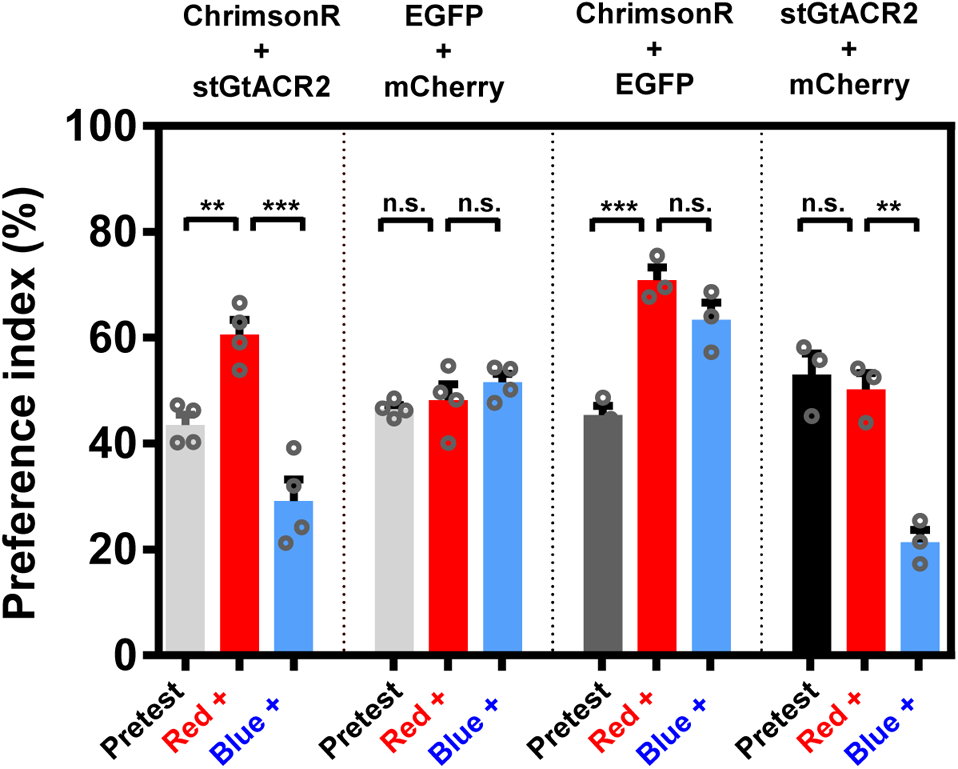
Summary of preference indices (the ratio of the time that mice spend in the left chamber to the whole recorded time) under red and blue stimulations for different mouse groups expressing ChrimsonR + stGtACR2 (*n* = 4 mice), EGFP + mCherry (*n* = 4 mice), ChrimsonR + EGFP (*n* = 3 mice), stGtACR2 + mCherry (*n* = 3 mice). Student’s *t* test, ** *P* < 0.01, *** *P* < 0.001, n.s. *P* > 0.05. The data for ChrimsonR + stGtACR2 and EGFP + mCherry groups are identical to those presented in Fig. 3f.

**Figure S21.**
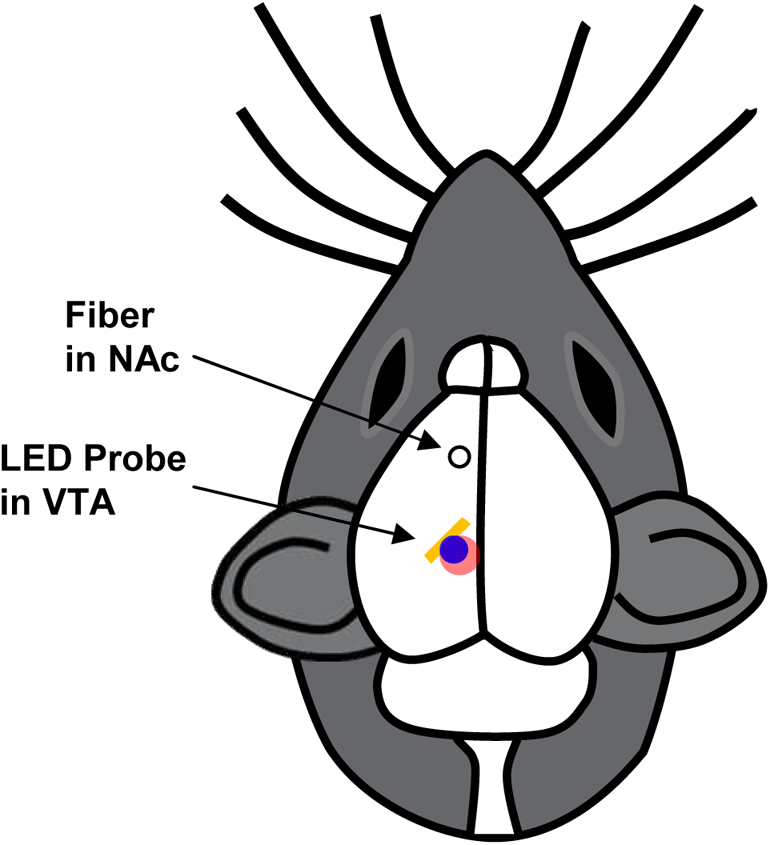
Schematic illustration (horizontal view) for the implantation of the fiber for DA recording in the NAc, and the dual-color LED probe for optogenetic stimulations in the VTA. The specific insertion angle is chosen for the LED probe, to minimize the optical crosstalk that disturbs the fiber photometric recording in the NAc.

**Figure S22.**
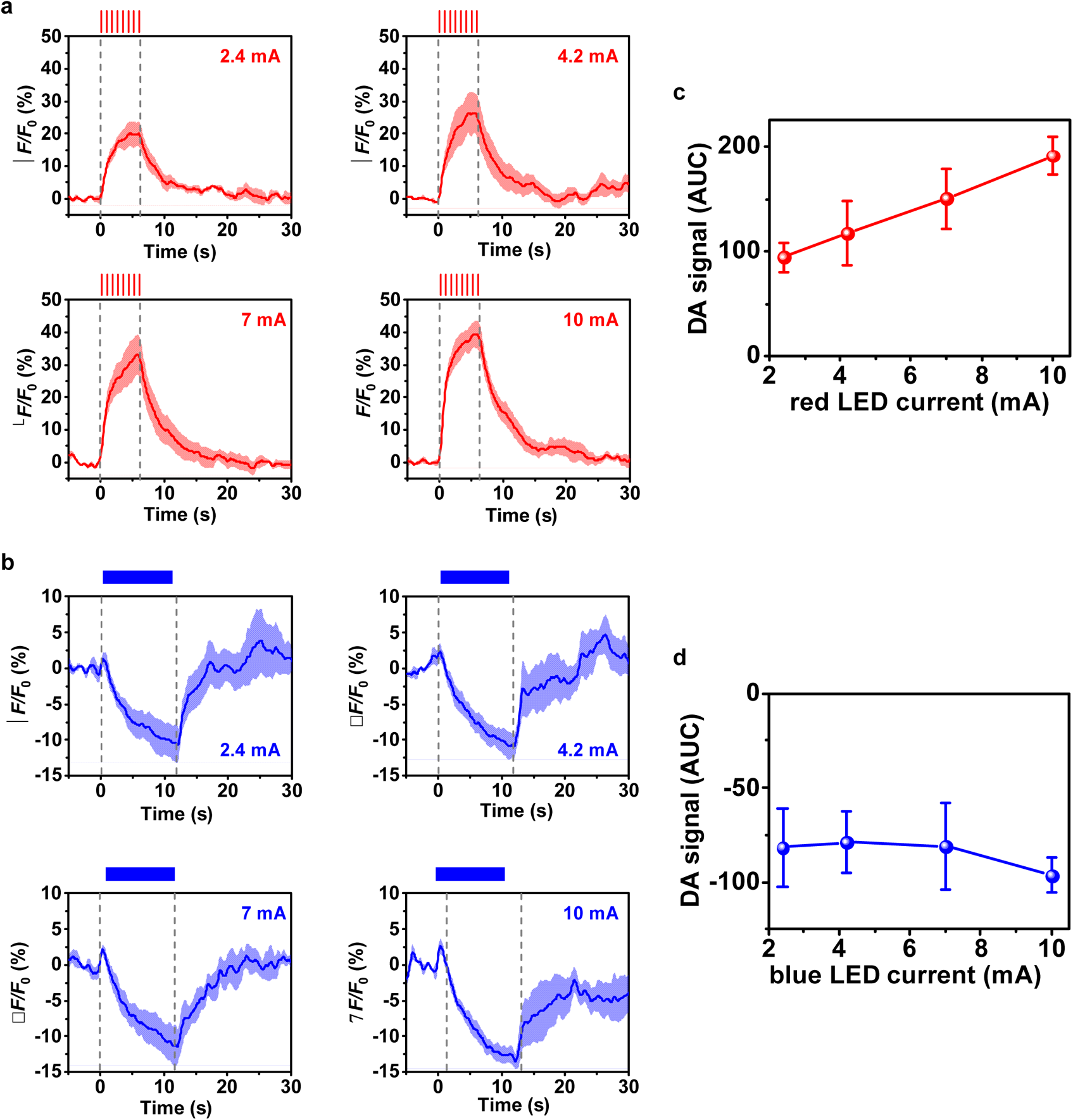
DA signals recorded as the fluorescence of GRAB_DA2m_ in the NAc, in response to optogenetic stimulations applied in the VTA by red or blue micro-LEDs. (a, b) Representative traces of DA signals in response to stimulations by (a) the red LED (20 Hz, 20 ms, 6 s) and (b) the blue LED (continuous, 12 s) at various currents. The solid lines and shaded areas indicate the mean and s.e.m., respectively. (*n* = 4 mice). (c, d) Accumulative DA signals in response to (c) red and (d) blue stimulations under different LED currents. AUC: area under the curve during stimulation. The data for red LED at 7 mA and blue LED at 2.4 mA, and (c), (d) are identical to those presented in Figure 4c–4e.

**Figure S23.**
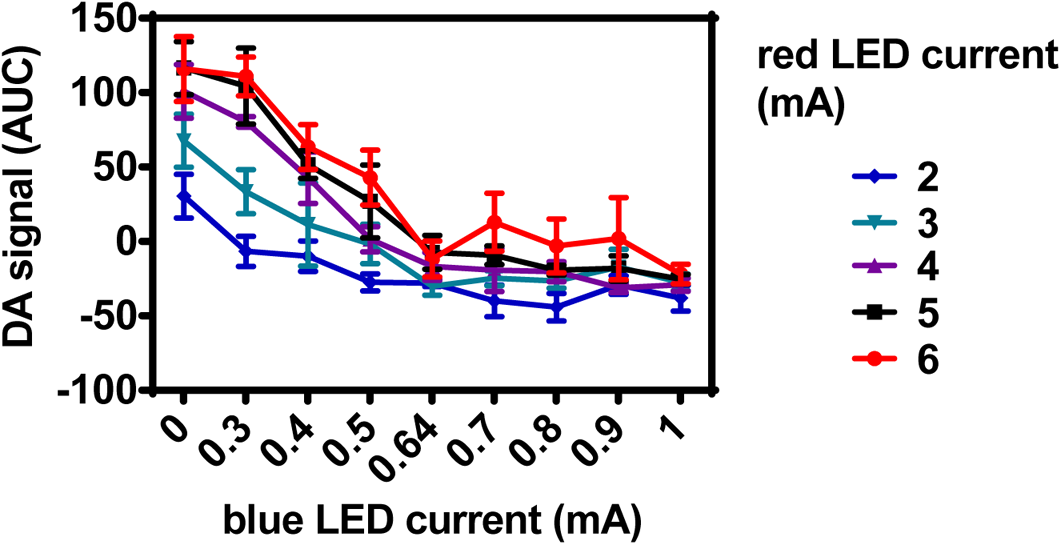
Detailed results of measured DA signals (AUC) with the combination of red and blue irradiance at different LED currents simultaneously for 6 s (*n* = 3 mice).

**Figure S24.**
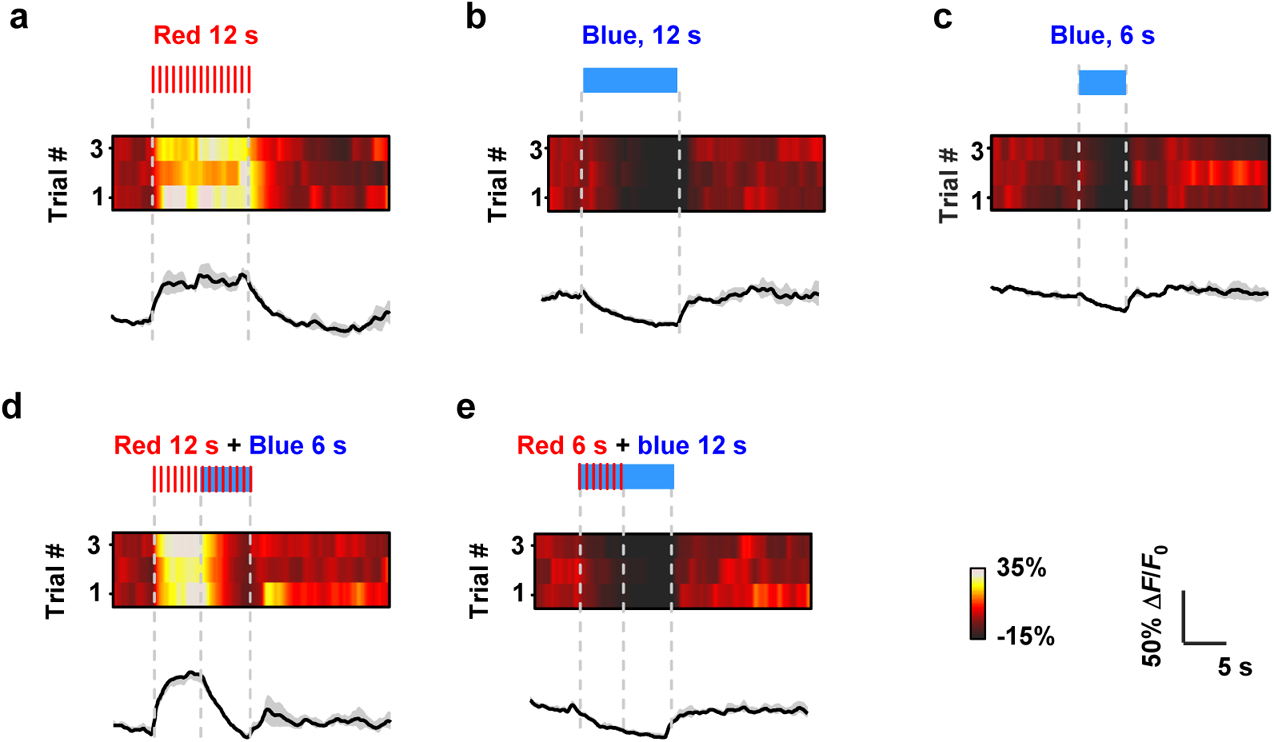
Example traces of optogenetically evoked DA transients for mice co-expressing stGtACR2 + ChrimsonR, in response to different stimulation patterns. (a) Red LED on, 20 Hz, 20-ms pulse, 10 mA, 0–12 s. (b) Blue LED on, continuous, 3 mA, 0–12 s. (c) Blue LED on, continuous, 3 mA, 6–12 s. (d) Red LED on 0–12 s and Blue LED on 6– 12 s. (e) Red LED on 0–6 s and Blue LED on 0–12 s. The solid lines and shaded areas indicate the mean and s.e.m., respectively.

**Movie S1.**
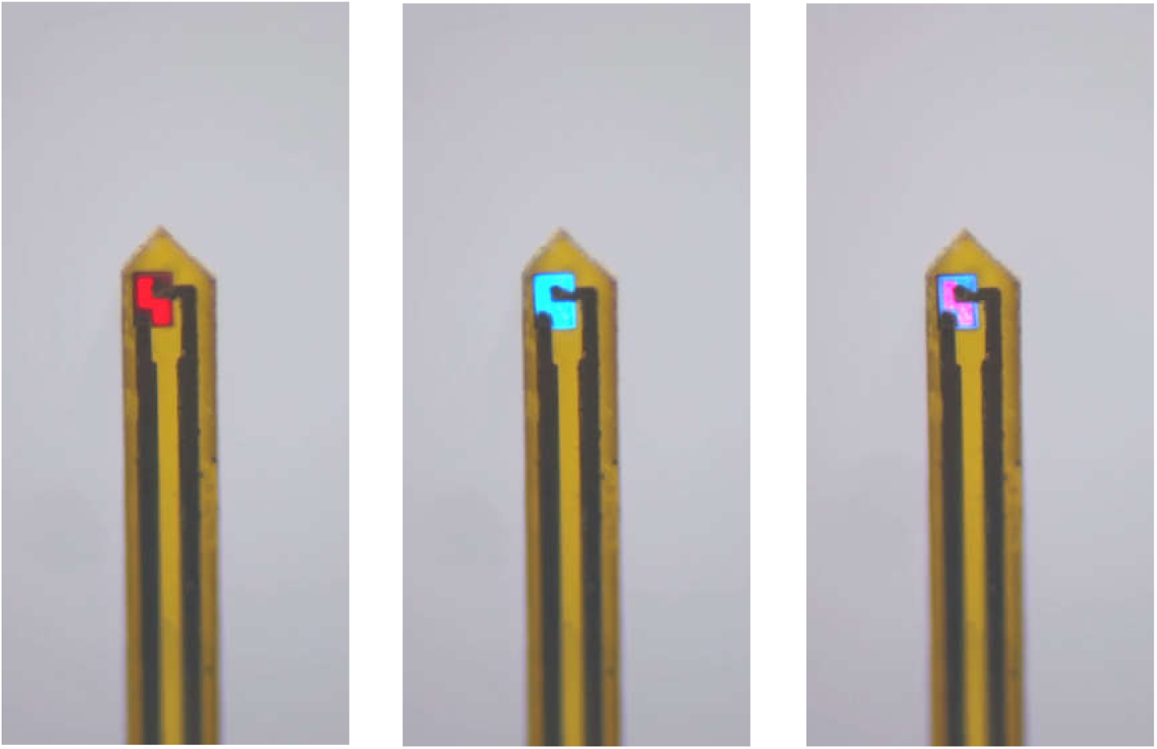
Video for dual-color micro-LED probe, displaying alternating blue and red emissions.

**Movie S2.**
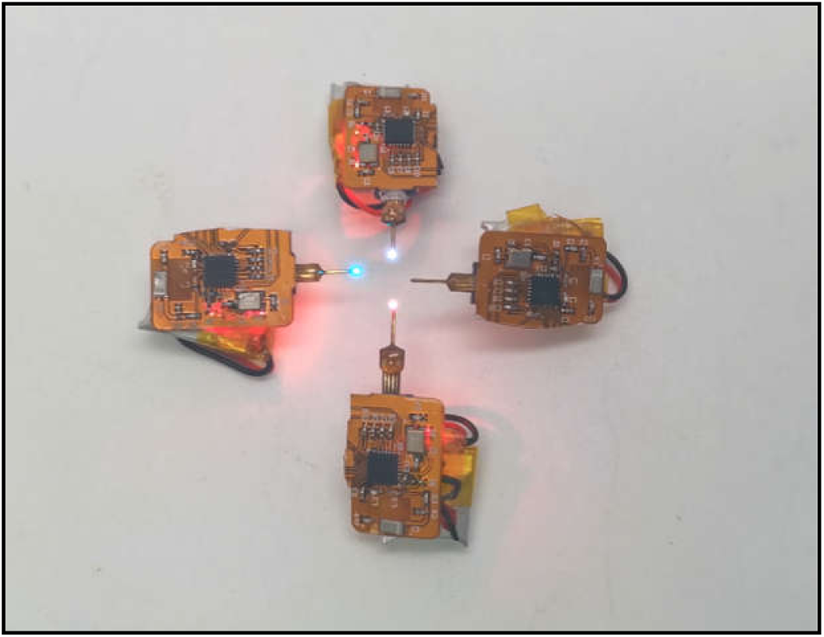
Video for multiple dual-color LED probes assembled with wirelessly operated circuit modules, showing capabilities for independent light emission control.

**Movie S3.**
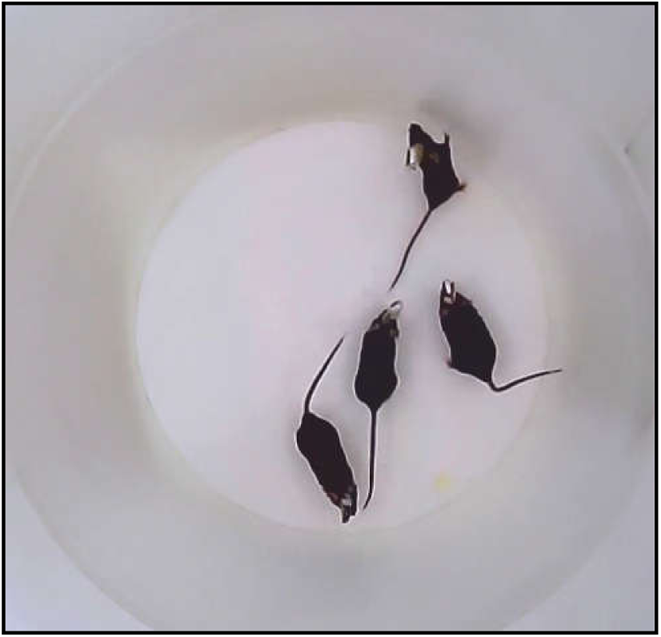
Video for four freely moving mice implanted with micro-LED probes and head-mounted circuits.

**Movie S4.**
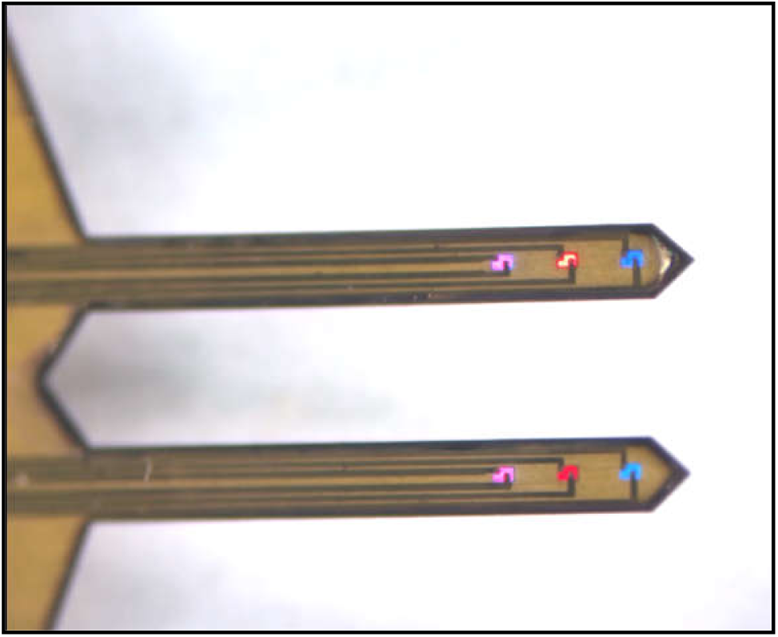
Video for multi-channel dual-color probes with 6 pairs of stacked red-blue micro-LEDs, displaying alternating blue and red emissions. These micro-LEDs are driven by a wired external power source.

**Movie S5.**
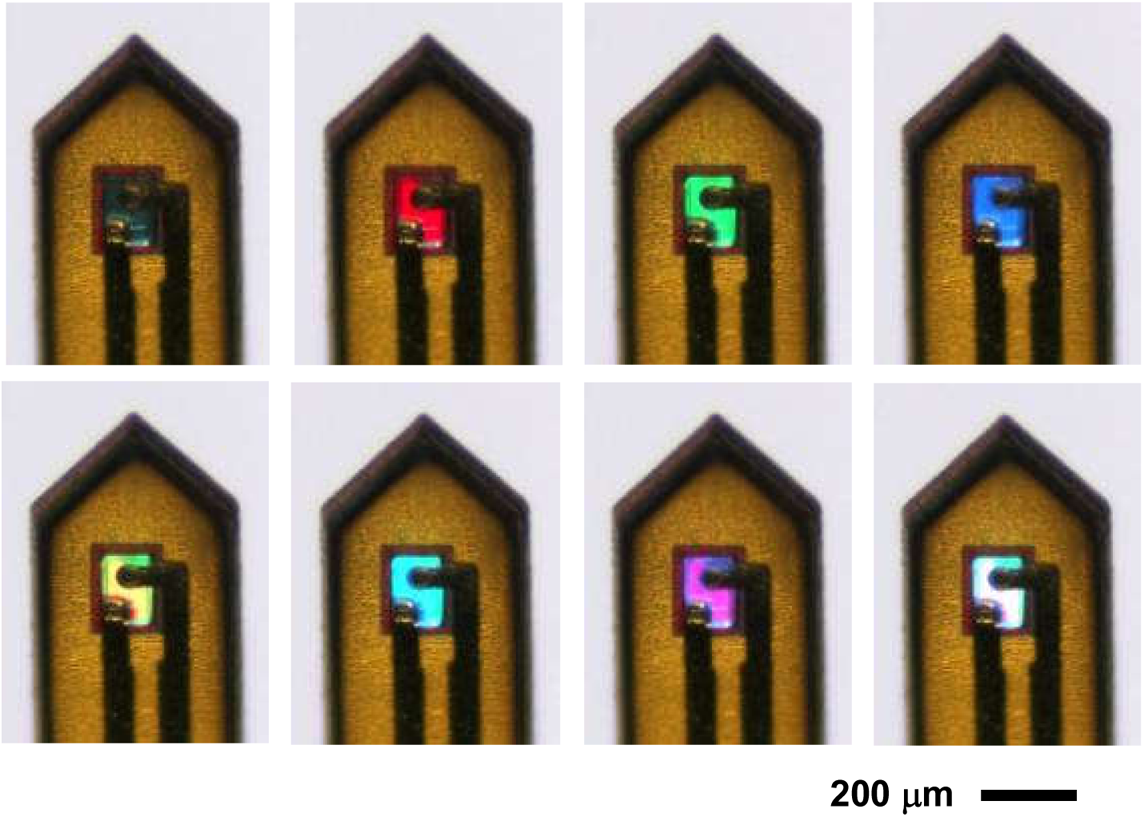
Video for a tri-color (red-green-blue) micro-LED needle structure, displaying different colors under current injection. These micro-LEDs are driven by a wired external power source.

